# Sampling Aware Ancestral State Inference

**DOI:** 10.1101/2025.05.20.655151

**Authors:** Yexuan Song, Ivan Gill, Ailene MacPherson, Caroline Colijn

## Abstract

In phylogeography, ancestral state inference methods are used to identify the geographic or host species origin of viral or bacterial lineages and reconstruct their transmission histories over time. However, differences in sampling among states can bias these inference methods. We introduce sampling-aware ancestral state inference (SAASI), a method that accounts for sampling differences. Here, we apply SAASI to the multi-host spread of the H5N1 virus in the United States in 2024 and find that the key transmission event from wild birds to cattle is estimated to occur later under lower sampling in wild birds (compared to other species) than when sampling is not accounted for. Using simulation, we find that SAASI infers past viral locations/host species considerably more accurately than standard methods when sampling bias exists, is computationally feasible for large datasets and scales to trees with 100,000 tips.

## Introduction

Ancestral state inference (sometimes called ancestral state reconstruction) refers to inferring the states of the ancestors of a set of taxa, given data about the states of the taxa. Depending on the state data used, ancestral state inference (ASI) has a broad range of applications. Given sequence data, it can be used in phylogenetics to infer the likely molecular sequences of ancestral organisms, for example, to estimate when and in which lineages a particular polymorphism (for example, conferring antibiotic resistance) emerged [1]. Alternatively, ASI is also used to infer traits such as a pathogen’s host species [2, 3], organisms’ geographic locations [4–6], accessory gene presence/absence [7, 8] or morphological or other traits [9, 10]. Phylogeography, in particular, is a major application of ancestral state inference inference, in which past geographic movements of viruses or other pathogens are reconstructed. For example, reconstructing the geographic spread of a virus can inform policy decisions about transportation and borders [11]. Understanding viral transmission between groups of a heterogeneous host population is important for reconstructing the origins of epidemics and designing effective control strategies. Estimates of interspecific transmission of zoonotic viruses can help identify the determinants of cross-species transmission events and, consequently, zoonotic risk.

Ancestral state inference methods use models that describe the process by which traits change over time, or by which the ‘state’ (e.g., geographic location, host species, molecular sequence, etc.) changes. Different traits or states may also be associated with different rates of speciation (branching), extinction (of the relevant lineage) and sampling. Taking this into account, the state-dependent speciation and extinction (SSE) family of models [12] can estimate trait-specific branching and extinction rates given a phylogenetic tree and traits of the sampled taxa. These include two states (BiSSE) models, multiple-state (MuSSE) models, hidden Markov (HiSSE) models, cladogenesis (ClaSSE) models and more [12–15]. Since trait-dependent speciation and extinction also impact the likelihood of a phylogenetic tree given sequence and trait data, these models inform phylogenetic reconstructions [16] in which state-dependent rates can be estimated alongside phylogenetic trees.

Inferring the states of ancestral organisms poses a distinct problem from estimating state-dependent rates. In ASI, each node in the phylogeny (and sometimes each point in the phylogeny) is associated with a particular state, or with a set of probabilities that the point is in each of the possible states. This may be done with stochastic character-mapping: mapping characters (or traits, or states) on to a phylogeny using a stochastic model [17–19], which is implemented for the case of a molecular evolution model in simmap [20]. Recently, Freyman and Höhna [21] developed a stochastic character mapping method for SSE models. This approach, however, is not yet widely used in viral (or pathogen) phylogeography. In this context, the states (e.g. geographic locations, host species) are often modelled as changing along a phylogenetic tree according to a continuous-time Markov chain, in a similar manner to how a molecular sequence evolves [22–24]: using a continuous-time Markov chain with a given instantaneous rate matrix *Q* specifying the rates at which the state changes [20, 22, 25–27]. Ancestral states are then inferred using post- and pre-order tree traversal algorithms that incorporate information from the child and parent nodes to infer the ancestral states. These are implemented, for example, in the ace function in R’s ‘ape’ package, and in PastML [26] and TreeTime mugration [27] model. A feature of viral geographic and/or host species data that is not yet represented in the SSE-based/stochastic mapping literature is the possibility of heavily biased sampling of lineages across space or among host species.

Sampling differences are known to impact phylogeographic estimates [4, 28, 28–31]. Down-sampling is often used to try to achieve relatively uniform sampling across locations through maximizing spatial or temporal coverage [11, 31–33], using epidemiological data as a reference point (e.g. hospitalizations vs cases) [34], or incorporating information about recent migration events and/or adding ‘sequence-free’ samples [5, 35]. Down-sampling, however, reduces the amount of data included in the analysis, may need to be replicated multiple times and is therefore time-consuming, and may not ultimately solve the problem. Furthermore, sampling differences may be extreme, as in the case of the recent influenza H5N1 outbreak, where samples from livestock and human cases are more readily available than those from wild animal populations. As genomic surveillance expands and we prepare for additional zoonotic spillover events that could lead to future large outbreaks or pandemics, it is essential to account for sampling bias and other state-dependent variation in ancestral state reconstruction on viral or other pathogen phylogenies.

In this work, we present sampling-aware ancestral state inference (SAASI), which builds on recent work on stochastic character mapping on fixed phylogenetic trees in state-dependent speciation and extinction (SSE) models [21]. We introduce two core developments: the inclusion of, and consequent adjustment for, sampling differences among states, including the sampling of viral lineages through time, and a modification to the stochastic character mapping method for SSE models that appropriately conditions on the observed tree. Our approach scales readily to very large phylogenetic trees. We test SAASI with simulated data, comparing it to ancestral character estimates from some widely used maximum likelihood methods: the ace function in R [36], simmap [20], PastML[26], and TreeTime[27]. As a proof of principle, we use SAASI to explore robustness to sampling for key host jumps and geographic movements in the recent avian influenza H5N1 outbreak in the United States [2].

## Results

SAASI is based on a 4-step algorithm that uses state-dependent speciation, extinction and sampling rates, solving differential equations along the edges of a tree, to assign ancestral state probabilities to internal nodes in the tree. The algorithm builds on our formalization of the stochastic mapping approach previously introduced [21], in which we condition on the part of the tree that is not descended from a focal node, and use the likelihood for the descending part. The algorithm uses a pre- and post-order tree traversal, as is common in such methods. The differential equations, conditioning and mathematical expressions are given in the Methods section.

To evaluate the performance of SAASI under sampling bias, we simulate trees using the birth-death-sampling process with two states, reconstruct ancestral states using SAASI and other methods, and compare the results. We illustrate the results first with one simulated tree, and then show results for a simulation study with a range of speciation and extinction rates, and a range of sampling biases, including unbiased sampling, and downsampling the data to achieve unbiased sampling (but with less data). We compare seven ancestral state inference methods: ace [22], PastML [26], simmap [20], TreeTime mugration model [27], TreeTime mugration model with sampling bias correction, SAASI with true parameters, and SAASI with estimated parameters (referred to as SAASI^*∗*^; we estimate the speciation, extinction rate, and transition rates). For each case we calculate two types of reconstruction accuracies: “consensus accuracy” gives the proportion of internal nodes where the consensus state (i.e., most likely ancestral state) matches the true state whereas “probability accuracy” is the average probability of the true state across internal nodes. Given that these two measures of accuracy broadly align, we discuss only consensus accuracy below with both measures reported in all tables and figures.

We apply SAASI to a timed phylogeny from the highly pathogenic avian influenza A (H5N1) outbreak in the US [2]. The phylogeny comprises 104 hemagglutinin (HA) segment sequences collected from April 2023 to April 2024, representing five host species (cattle, mammals, poultry, and wild birds) across 12 US states. Our analysis is a proof of principle for SAASI, rather than a re-analysis of the full dataset (which has over 15,000 sequences). We examine the effect of accounting for lower sampling in wild bird populations than in domestic animals and humans, using the host type as the state. To illustrate the various applications of SAASI, we also explore the geographic spread of H5N1, accounting for the possibility of higher sampling in Texas than other states [37, 38].

Detailed descriptions of the simulation parameters, method settings, accuracy calculations, and H5N1 analysis are described in the Methods section.

### Simulation study

Figure 1 shows SAASI’s performance on an example simulated tree for a binary state model with highly biased sampling (specifically, state 1 is sampled at one-tenth the rate of state 2), but where the states have no other effects on diversification. When the true sampling rates are known, SAASI correctly infers most internal nodes with consensus accuracy *a*_consensus_ = 0.98. In contrast, ace incorrectly infers approximately half of the internal node states (*a*_consensus_ *≈* 0.5) and misidentifies the root state as state 2. Under uniform sampling assumptions (Supplementary Fig. 1), SAASI’s reconstruction is very similar to that of ace and simmap.

**Figure 1.**
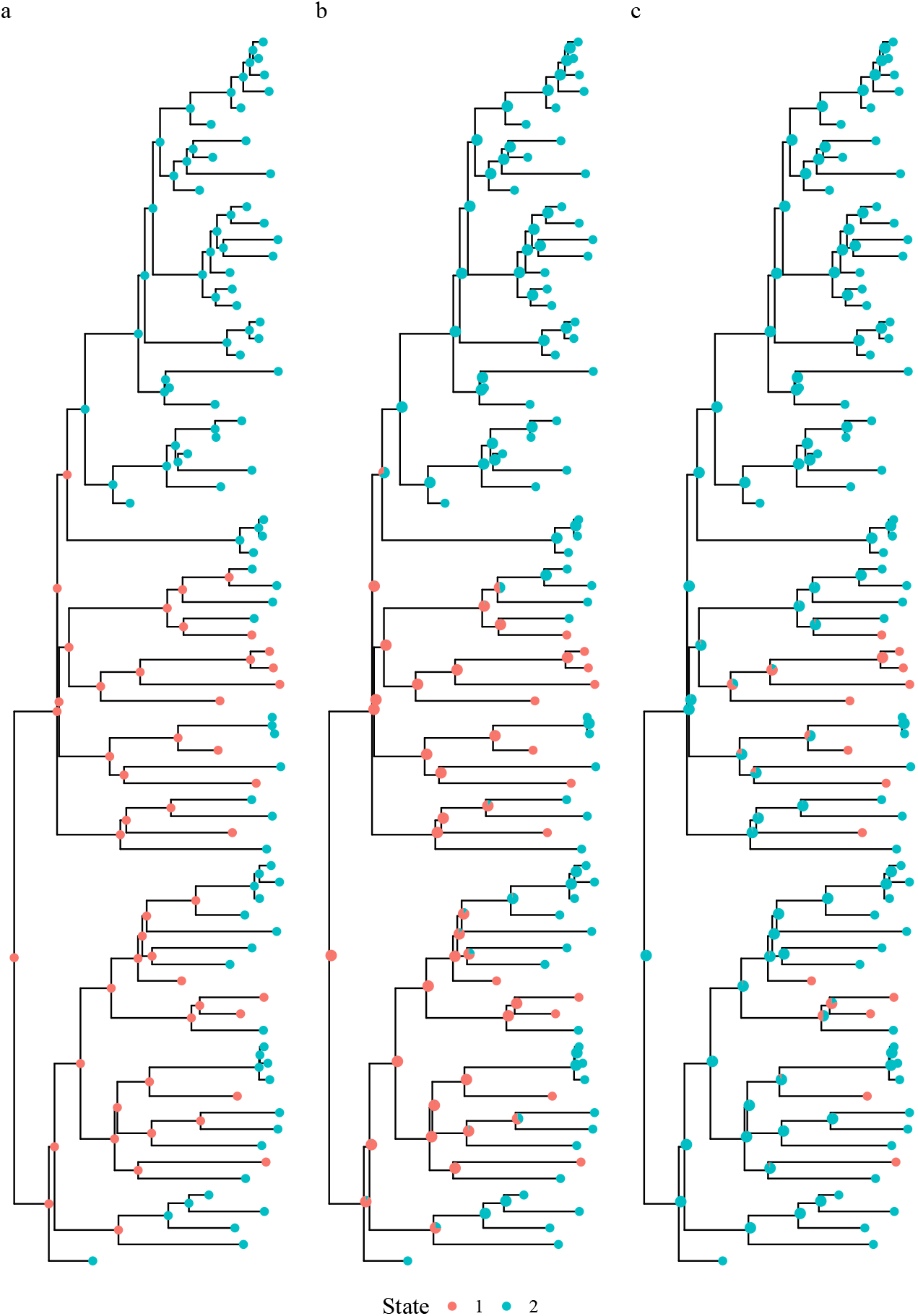
Ancestral state inference using SAASI and ace. a: Simulated tree with known transmission histories; b: SAASI with true parameter values; c: ace with ‘ER’ transition model. Pie charts indicate the inferred probabilities of being in particular states. The tree was generated with the following parameters: *λ*_1_ = *λ*_2_ = 1, *µ*_1_ = *µ*_2_ = 0.045, *q*_12_ = *q*_21_ = 0.05, and *ψ*_1_ = 0.05, *ψ*_2_ = 0.5.

Complementing this single example tree analysis, we compare SAASI to other ancestral state inference methods (ace, PastML, simmap, TreeTime, Treetime with sampling correction) across 1,000 simulated trees under high, moderate, and no sampling bias (Figure 2; see Supplementary Figs. 6, 9). Under high sampling bias (*ψ*_2_ = 10*ψ*_1_, mean tree size 463 tips), SAASI substantially outperforms existing methods (*a*_consensus_ = 0.934 *±* 0.021). SAASI^*∗*^ also has high accuracy (*a*_consensus_ = 0.920 *±* 0.023). TreeTime with sampling bias correction has accuracy (*a*_consensus_ = 0.699 *±* 0.147). Under moderate sampling bias (*ψ*_2_ = 4*ψ*_1_, mean tree size 2669 tips), all methods have high accuracy. If sampling is uniform across all states (*ψ*_1_ = *ψ*_2_), all methods have high accuracy; ace, simmap, and TreeTime all have consensus accuracies of *a*_consensus_ = 0.98. SAASI obtains slightly lower consensus accuracy *a*_consensus_ = 0.977 *±* 0.008, SAASI^*∗*^ obtains *a*_consensus_ = 0.973 *±* 0.010, and PastML obtains *a*_consensus_ = 0.979 *±* 0.010.

**Figure 2.**
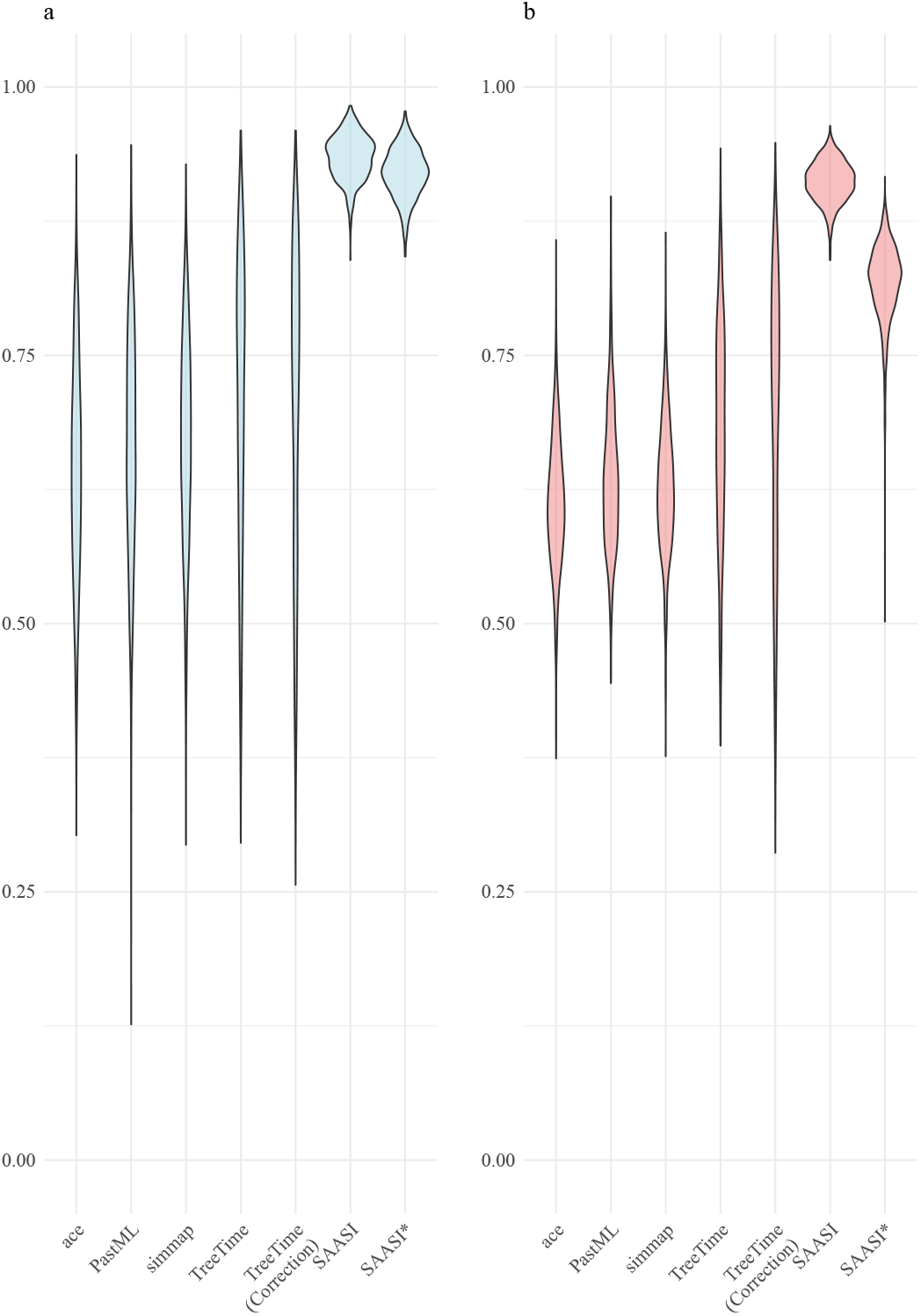
Accuracy of ancestral state reconstruction methods under highly biased sampling (*ψ*_2_ = 10*ψ*_1_) and state dependent diversification across 1000 simulated trees. a: Consensus accuracy, defined as the fraction of correctly inferred ancestral node states. b: Probability accuracy, accounting for uncertainty in the node inference. Violin plots show the distribution of accuracy values across all simulations. ace and simmap use the equal-rates (ER) transition model. PastML employs the MPPA method with the F81 evolutionary model. TreeTime uses the mugration model with and without sampling bias correction (shown in parentheses). SAASI uses the true parameter values that generated the tree. SAASI^*∗*^ uses estimated speciation and extinction parameters, but is provided with the true sampling rates. Trees were generated with speciation rates *λ*_1_ = 3 and *λ*_2_ = 1.5, extinction rates *µ*_1_ = 0.1 and *µ*_2_ = 0.05, transition rates *q*_12_ = *q*_21_ = 0.3, and sampling rates *ψ*_1_ = 0.1 and *ψ*_2_ = 1.0.

Supplementary Fig. 7 and Supplementary Table 3 explore the practice of downsampling data to obtain approximately uniform sampling. Under high sampling bias, downsampling achieves high accuracy when evaluated only on preserved nodes (red violin plots in Supplementary Fig. 7). However, if we account for a lack of inference on all internal nodes that are removed (green violin plots in Supplementary Fig. 7), the accuracy drops dramatically, and is substantially lower than reconstructions on the full tree (blue violin plots in Supplementary Fig. 7). Similar patterns arise under moderate sampling bias (Supplementary Fig. 8 and Supplementary Table 4, where downsampling reduces accuracy when missing transition events are considered.

When the sampling ratio is mis-specified in SAASI (Supplementary Fig. 2), both SAASI’s consensus and probability accuracy decrease as the sampling ratio deviates from the true sampling ratio (here 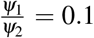 to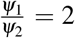). If the sampling ratios are close to the true sampling ratio (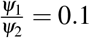 to 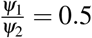), the accuracy exceeds 0.75. However, if the sampling ratio is mis-specified (such that 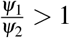), the accuracy is reduced to approximately 0.5, comparable to that of reconstructions using ace. In addition to assessing performance under the true parameters (SAASI) and inferred state transition rates (SAASI^*∗*^), we similarly considered the sensitivity of our method to mis-specified transition rates (*Q*), as shown in Supplementary Fig. 3. We find that inference is insensitive to this mis-specification.

### Scalability to larger trees

SAASI has a similar running time to simmap, with a mean of 25 seconds, and average tree sizes between 1,100 and 1,600 tips. In contrast, ace, PastML and TreeTime both have fast completion time, with an average of 1 second for ace and 5 seconds for PastML and TreeTime (Supplementary Fig. 10). SAASI is scalable for trees that contain more than 100,000 tips. We estimate that the relationship between tree size and running time (in seconds) is linear. For trees with 100,000 tips, SAASI completes in approximately 1 hour (3500 seconds; Supplementary Fig. 11). The running time scales quadratically with the number of states *k* (Supplementary Fig. 12). The overall computational complexity is *𝒪* (*nk*^2^), where *n* is the number of internal nodes in the tree and *k* is the number of states.

Our simulations indicate that SAASI has strong performance under a range of sampling biases, parameter settings, and parameter choices, and is robust to some mis-specification of its input parameters. SAASI is feasible for very large trees, for example running in under an hour for a tree with 100K tips and two states. SAASI, of course, does not reconstruct the phylogenetic tree itself, but runs on an input tree (and it would be feasible to run it on a collection of trees, such as a posterior sample). SAASI is most appropriate for situations where there is known, extensive sampling bias (the rates need not be known precisely); where the dataset is relatively large (prohibiting Bayesian phylogenetics) and where ancestral state inference is the aim (as opposed, for example, to rate estimates). Figure 3 has a decision tree to guide the choice of ancestral state inference tools.

**Figure 3.**
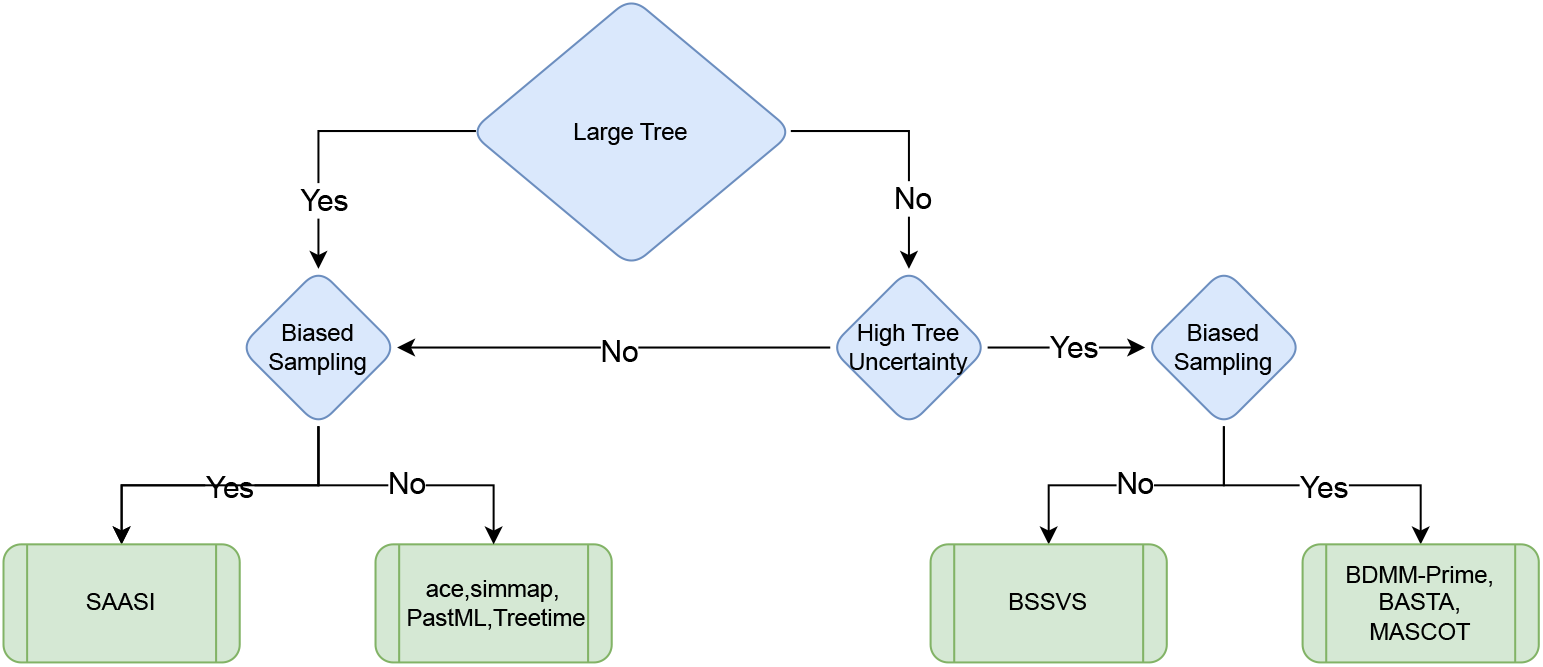
Decision tree for Ancestral State Inference method selection. The flowchart guides method selection (green rectangles) based on the characteristics of the data and sampling process (blue diamonds). Specifically, SAASI is suitable for large phylogenetic trees with biased sampling. When sampling is uniform, maximum-likelihood methods (ace, simmap, PastML, TreeTime) are appropriate. Bayesian methods (BSSVS, BASTA, MASCOT, BDMM-Prime) accommodate phylogenetic uncertainty (tree topology and branch lengths) but are limited to smaller datasets and typically require longer run times.

### Avian influenza

Adjusting for lower sampling of wild bird infections than infections in other hosts, we find that there may have been more than one transmission of H5N1 from wild birds to cattle (Figure 4). We infer 10 transmissions from wild birds to cattle when wild birds are sampled at 1/10 the sampling rate of other states, and 12 transmissions when wild birds are sampled at 1/100 the others’ sampling rate. These are inferred to have occurred from late January to early February 2024 (Figure 4 b,c, between red dashed lines), slightly later than the previous estimate [2]. In contrast, if we model no sampling difference between species, both ace and SAASI cannot identify the key transmission events from wild birds to cattle (Supplementary Fig. 4).

**Figure 4.**
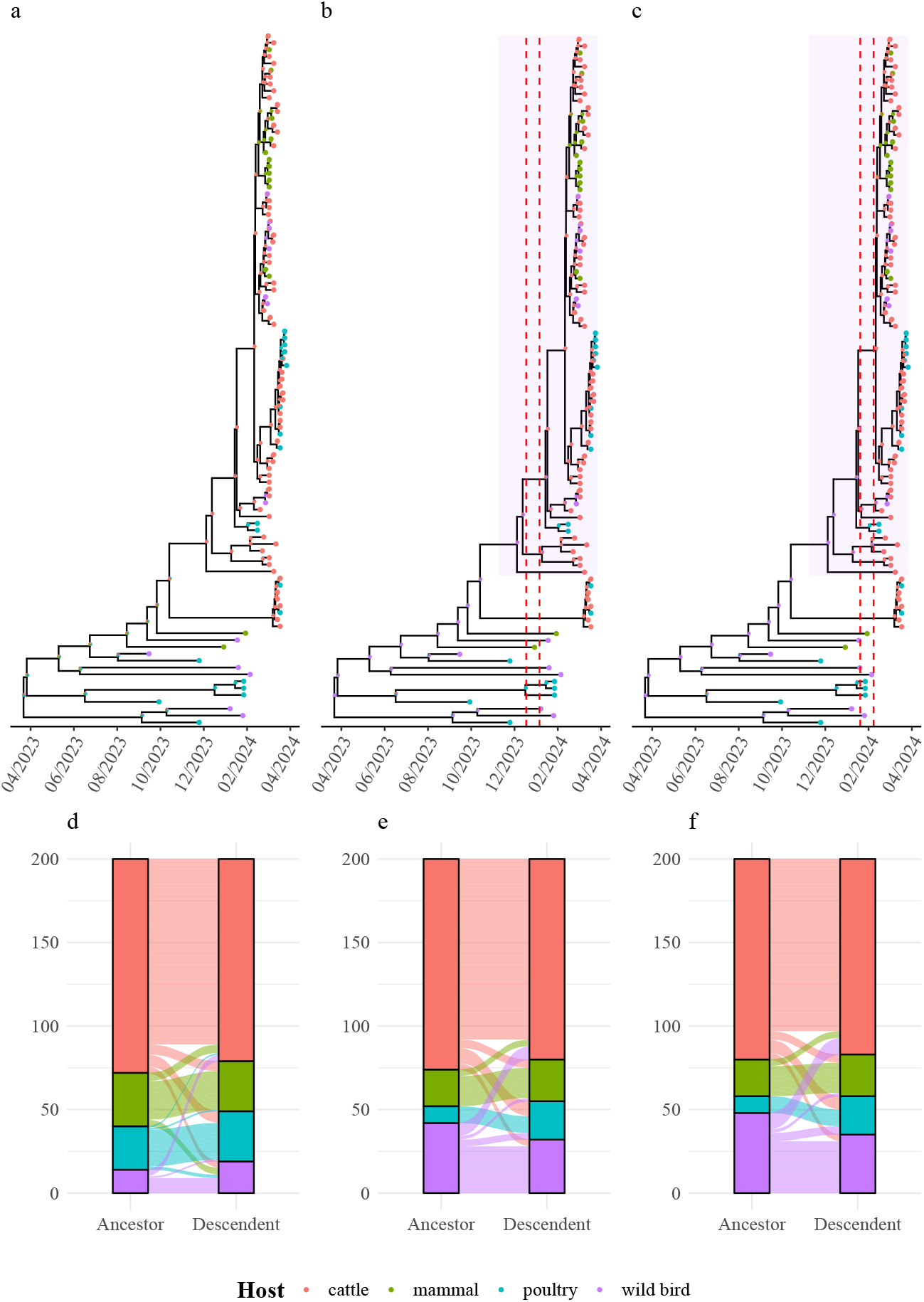
Ancestral state inference of the H5N1 HA segment tree using SAASI under different host-level sampling models. a: Inferred hosts under equal sampling rates; b: Wild birds at one-tenth sampling; c: Wild birds at one-one hundredth sampling; d: Inferred viral transitions between hosts in a; e: Inferred viral transitions between hosts in b; f: Inferred viral transitions between hosts in c. Pie charts indicate the inferred probabilities of being in particular states. Transition rates of the wild bird population are adjusted by a factor of 2 (*c* = 2). The dashed red lines indicate the key transition event from wild bird to cattle.

We analysed viral movement patterns among host taxa between parent and child node pairs in the phylogeny, and illustrated these with alluvial plots (Figure 4 panels d–f). We identify the internal node as being in state *i* if *i* has the highest probability (i.e., the consensus state). This allows us to probe not just where a particular node was located, but to ask about the nature of viral movements between taxonomic groups. We find that cattle are the origin species for most host species jumps, under all sampling models we explored. We find that adjusting for under-sampling of wild bird infections leads to the inference that more nodes of the phylogeny are in wild birds, compared to scenarios with more uniform sampling. This, in turn, means fewer inferred transitions from other populations into wild birds (see Figure 4 d compared to e and f). In particular, without accounting for sampling, standard methods infer transitions from both other mammals and poultry into wild birds (as well as cattle). With sampling adjustment, we would infer that only cattle infections were ever reintroduced to wild birds. We also find more transitions from wild birds into other hosts when we account for lower sampling in wild birds (i.e. from wild birds to mammals and to poultry, along with cattle). These results are robust to the specific transition rate adjustment (Supplementary Fig. 5).

Figure 5 shows the results of our H5N1 geographic analysis in which we examined the inferred geographic origin of the clade predominantly in cattle. If we model that there is no sampling difference between jurisdictions (left) compared to modelling a factor of 5 sampling difference between Texas and other states (right) in Figure 5, both models suggest that the first spillover event from the wild bird population to cattle occurred in Texas. If we model that there is no sampling difference between states, SAASI would infer that it is highly likely that a wild bird moved from Texas to New Mexico, leading to the emergence of a predominantly New Mexico clade in March 2024. In contrast, if we model that infections in Texas were sampled at a rate 5 times higher than in other states, SAASI would suggest that there was a wild bird movement from New Mexico to Texas between November 2023 and January 2024 (red dashed lines). In either case, our analyses show that the virus rapidly spread from Texas to other states in March 2024, including Ohio, Kansas, Michigan, and New Mexico. Overall, the geographic reconstructions are robust to a five times sampling difference between Texas and other states. Phylogeographic analysis in [2] also inferred significant interstate movement, with confirmed transitions from Texas to Kansas, New Mexico, and Michigan. Epidemiological records also documented the movements of infected cattle from a Texas herd to North Carolina and Idaho [39].

**Figure 5.**
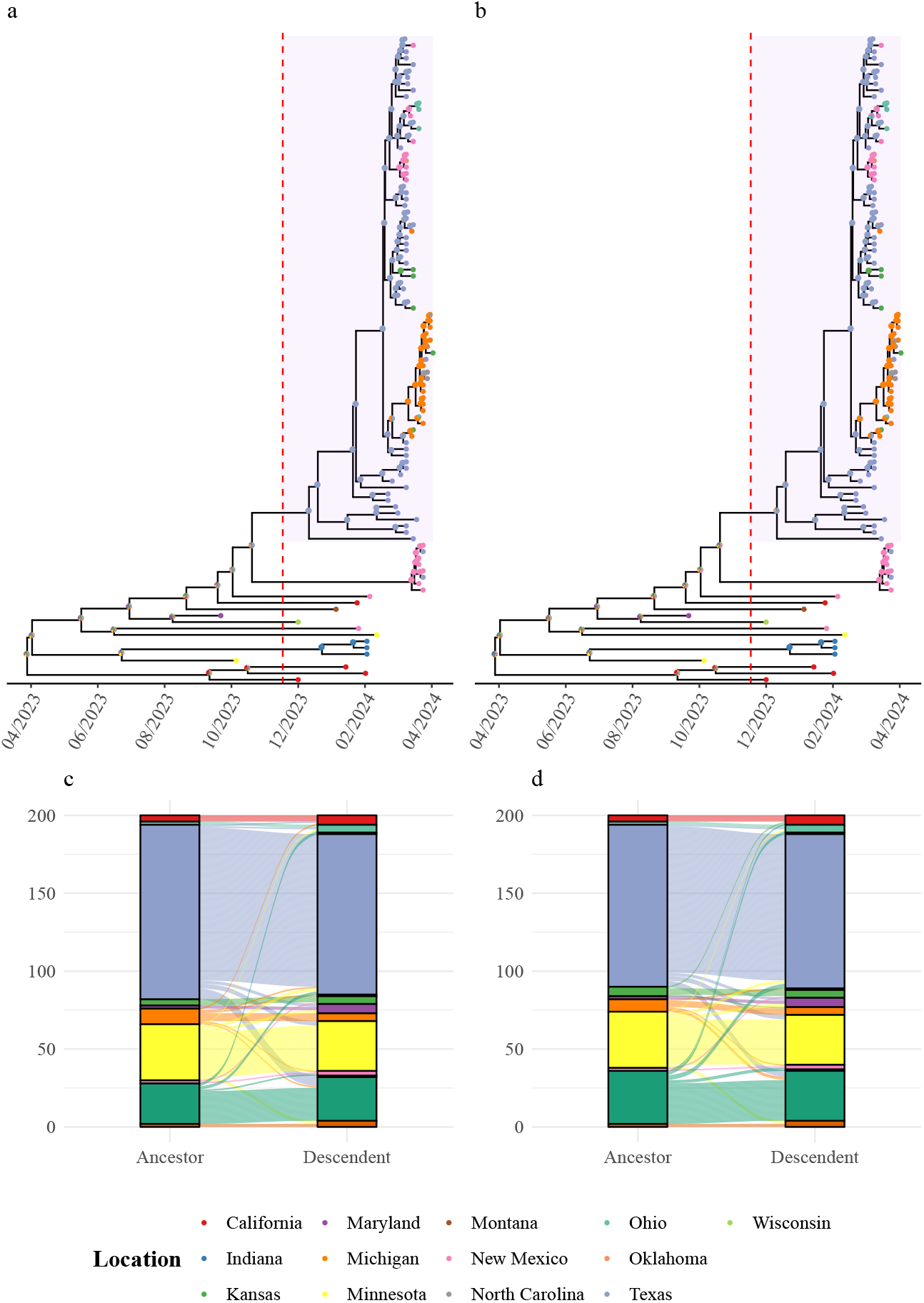
Phylogeographic ancestral state inference using SAASI under different sampling models for Texas and other US states. a: Inferred geographic locations under equal sampling; b: Texas at five times sampling. c: Inferred viral transitions between geographic locations in a; d: Inferred viral transitions between geographic locations in b. Pie charts indicate the inferred probabilities of being in ancestral states. The transition rates between states are modelled as equal and estimated using ace. The dashed red line indicates the estimated time it takes for the virus to move to Texas.

## Discussion

We have introduced and implemented a fast approach for sampling-aware ancestral state inference, SAASI, which applies to a range of viral phylogeography applications, including the reconstruction of ancestral host species or geographic locations in large phylogenies. A key methodological contribution of SAASI is a modification to the pre-order traversal algorithm used. Unlike standard approaches, we exclude birth and extinction events in our pre-order traversal, because we are conditioning on the observed tree topology not descending from the focal node. In simulations, SAASI achieves a high accuracy in the presence of sampling bias, in contrast to standard methods, which are negatively impacted by sampling differences among states. SAASI scales readily to very large phylogenies. SAASI removes the need to downsample data to achieve near-uniform sampling, a common practice. This is of particular help for highly biased sampling, where achieving uniformity by downsampling requires removing a large portion of the data, with all the associated limitations.

As a proof of principle, we applied SAASI to a small dataset from the recent outbreak of avian influenza H5N1, using 104 HA sequences from six host species in 12 geographic locations to demonstrate how sampling assumptions affect phylogeographic inference. Nguyen et al. [2] conducted a comprehensive analysis using the complete H5N1 phylogeny across multiple viral segments (HA, PB2, PA, NP, NA, MP, and NS). We find that SAASI is able to identify the time of transition from wild birds to cattle (from late January to early February 2024), and infers that the root of the clade that was mainly in cattle was likely to have been in wild birds, with several subsequent introductions to cattle. These findings emphasize the importance of accounting for sampling bias, which significantly impacts phylogeographic reconstruction. However, drawing conclusions about the actual number of spillover events would require a comprehensive re-analysis of the H5N1 data, which is beyond the scope of this work.

SAASI has important limitations. First, the sampling difference under consideration is modelled as known (if it is not known, then SAASI offers an approach to sensitivity analysis). If attempts were made to estimate sampling bias alongside transition rates, state-dependent branching and death rates, and states of internal nodes, it is likely that these would be collectively identifiable [40]. For example, having fewer taxa with a trait (and fewer branching events with an apparent lower rate) could be a result of either a lower branching rate or lower sampling intensity. Other natural trade-offs would affect the simultaneous estimation of the complete set of parameters. Furthermore, SAASI does not (yet) estimate other relevant parameters; in our application, these were estimated separately based on either birth-death models or ace’s estimates with appropriate adjustments. These could, in principle, be estimated in a first pass using one of the BiSSE [12] family of models without ancestral state reconstruction, but at present these require ultrametric trees and so are not suited for longitudinally sampled pathogen datasets.

We have focused on sampling differences, as it is likely to vary by location and host species, and this is a recognized challenge for phylogeographic reconstructions [28]. We note that evolutionary parameters, such as the molecular clock, substitution model, as well as branching and extinction rates, are likely to change from host to host, and rapid adaptation with selection may occur immediately following a host jump. These phenomena are challenging to include in phylogenetic inference. We used a fixed, timed phylogenetic tree because we aim to account for sampling differences in phylogeographic reconstructions at large scales.

In recent work, Vaughan and Stadler [40] also build upon the previous work of Freyman and Höhna [21], in their case to develop Bayesian inference for multi-type population trajectories. Their algorithms jointly infer the phylogenetic tree, multi-type birth-death model parameters (including our state-dependent branching, death, and sampling rates), ancestral node states, and type-specific population trajectories. Potentially unidentifiable combinations can be managed in Bayesian analyses (through priors, and through sampling a posterior, which may have complex correlation structure and strong correlations, but which reflects the estimated uncertainty). The sampling is done with a combination of particle filtering and Markov Chain Monte Carlo (MCMC) sampling. Other Bayesian phylogeography tools include discrete trait models with Bayesian stochastic search variable selection (BSSVS) [5, 41] (but without adjustment for differential sampling) and structured coalescent models [42, 43], which can accommodate sampling differences but which are highly computationally demanding and make coalescent assumptions for the demes. None of these approaches are feasible for trees with thousands of taxa or more.

In contrast to Bayesian approaches, SAASI can operate at very large scales and provide rapid estimates of ancestral node states, accounting for known sampling differences. This focus is motivated in part by the emergence of large-scale genomic surveillance efforts and phylogeographic analyses, in particular for SARS-CoV-2 [32, 44] but also for a wide range of pathogens and organisms, including influenza viruses [32, 45, 46] and other pathogens [47, 48]. In simulations, SAASI obtains a high accuracy and will be broadly relevant, as testing and sequencing policies vary from one jurisdiction to another. Despite advances in Bayesian methods, the current state of the art in phylogeography at large scales is to down-sample the data to approximately obtain representative sampling, and re-run analyses using tools like ace [33, 49, 50]. Repeat analyses may be time-consuming, give variable results, and always require discarding data. This approach also makes strong assumptions, in particular that the traits (here, host species or locations) do not affect the branching, sampling or death rates.

SAASI is less restrictive. It offers quick, robust and principled phylogeographic reconstructions that account for sampling bias. SAASI is suited for large phylogenetic trees with known or suspected sampling bias, making it applicable to scenarios where rapid analysis of large-scale surveillance data is required, and to ancestral state inference problems in any other large datasets. SAASI is relatively robust to parameter mis-specification, at least as explored in our simulation study, and is quick enough that determining the sensitivity to uncertain rates and phylogenetic uncertainty is feasible.

## Methods

We consider a rooted binary phylogenetic tree 𝒯 with known tree topology and character states (e.g. location, host species identification) at the tips. We assume that this tree is the result of a state-dependent birth-death-sampling process allowing for both cladogeneic and anagenic state changes. Specifically, a lineage in state *i* gives birth (e.g., viral transmission in the epidemiological context) to daughter lineages in states *j* and *k* at rate *λ*_*ijk*_ which define the 3D array Λ, dies (e.g., host recovery) at rate *µ*_*i*_ which define the vector 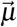, and transitions to state *j* at rate *q*_*ij*_ which define the matrix *Q* (see Table 1 for a list of notation). In addition to these rates, we introduce the state-dependent sampling rate 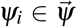 through time, with which viral sequences are collected throughout an ongoing epidemic. Finally, let *π*_*i*_ be the prior probability that the root *R* of the phylogenetic tree is in state *i* (defining the vector 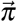).

**Table 1.**
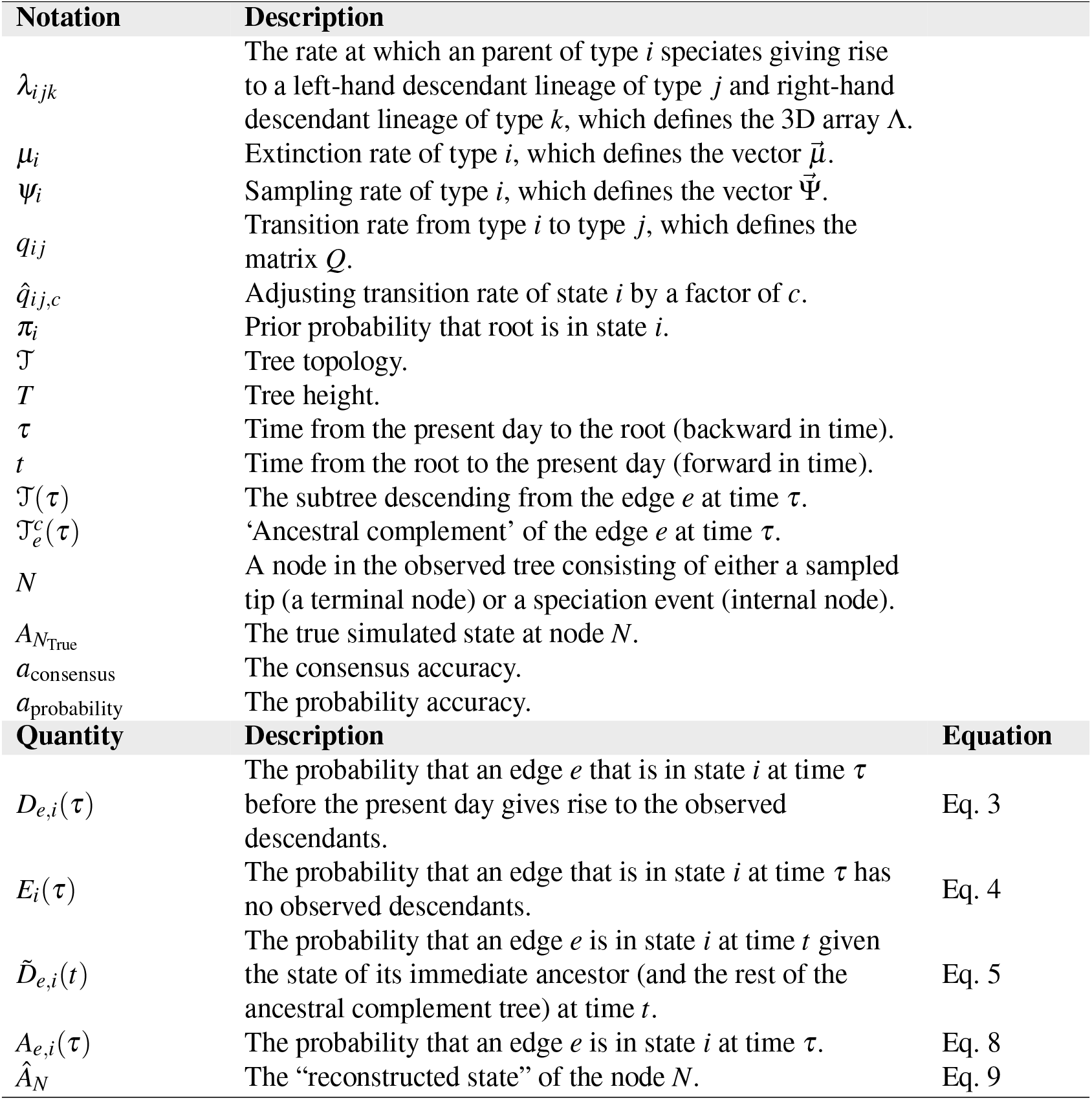
List of notation, derived quantities and corresponding equations. Throughout, time *τ* is measured backwards in time from the present day (*τ* = 0) to the root (*τ* = *T*) of the tree. Note that the tree 𝒯 includes information on the states at the tips of the tree. Similarly, 𝒯_*e*_ includes the tip states of the descendants of edge *e* and the ancestral complement 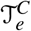 includes information on the states at all the tips in the tree except those that descend from edge *e*.

| Notation | Description |  |
| --- | --- | --- |
| $\lambda_{ijk}$ | The rate at which an parent of type $i$ speciates giving rise to a left-hand descendant lineage of type $j$ and right-hand descendant lineage of type $k$, which defines the 3D array $\Lambda$. | |
| $\mu_i$ | Extinction rate of type $i$, which defines the vector $\vec{\mu}$. | |
| $\psi_i$ | Sampling rate of type $i$, which defines the vector $\vec{\Psi}$. | |
| $q_{ij}$ | Transition rate from type $i$ to type $j$, which defines the matrix $Q$. | |
| $\hat{q}_{ij,c}$ | Adjusting transition rate of state $i$ by a factor of $c$. | |
| $\pi_i$ | Prior probability that root is in state $i$. | |
| $\mathcal{T}$ | Tree topology. | |
| $T$ | Tree height. | |
| $\tau$ | Time from the present day to the root (backward in time). | |
| $t$ | Time from the root to the present day (forward in time). | |
| $\mathcal{T}(\tau)$ | The subtree descending from the edge $e$ at time $\tau$. | |
| $\mathcal{T}_e^c(\tau)$ | ‘Ancestral complement’ of the edge $e$ at time $\tau$. | |
| $N$ | A node in the observed tree consisting of either a sampled tip (a terminal node) or a speciation event (internal node). | |
| $A_{N_{\text{True}}}$ | The true simulated state at node $N$. | |
| $a_{\text{consensus}}$ | The consensus accuracy. | |
| $a_{\text{probability}}$ | The probability accuracy. | |
| Quantity | Description | Equation |
| $D_{e,i}(\tau)$ | The probability that an edge $e$ that is in state $i$ at time $\tau$ before the present day gives rise to the observed descendants. | Eq. 3 |
| $E_i(\tau)$ | The probability that an edge that is in state $i$ at time $\tau$ has no observed descendants. | Eq. 4 |
| $\tilde{D}_{e,i}(t)$ | The probability that an edge $e$ is in state $i$ at time $t$ given the state of its immediate ancestor (and the rest of the ancestral complement tree) at time $t$. | Eq. 5 |
| $A_{e,i}(\tau)$ | The probability that an edge $e$ is in state $i$ at time $\tau$. | Eq. 8 |
| $\hat{A}_N$ | The “reconstructed state” of the node $N$. | Eq. 9 |

Here, we propose and implement a method for calculating the probability that at any time *τ* before the present day, an edge *e* in the phylogeny was in state *i ∈ S*, denoted *Y*_*e*_(*τ*) = *i*, accounting for both the observed states at the tips of the tree and the topology of the tree resulting from state-dependent diversification. Using the notation above, we wish to calculate:

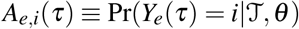

where 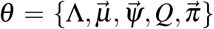 is the diversification model. Throughout, we define this kind of shorthand notation for each key probability, e.g. *A*_*e,i*_(*τ*). To calculate this focal probability, we formalize and extend the stochastic-mapping method presented by [21]. Our method involves first a post-order traversal of the tree (from tips to root) followed by a pre-order traversal (from root to tips) during which the ancestral state probabilities are obtained. To avoid confusion on the direction of time, throughout, we use *T* to denote the height of the tree, *τ* to denote time from the present day to the root of the tree, and *t* to denote time from the root to the present day such that *τ* = *T − t*. In line with SSE models, for functions that are neither more naturally described backward vs. forward in time (e.g., *A*_*e,i*_(*τ*)), we will use backward-in-time notation. We give an overview, rather than a detailed derivation, of the pre- and post-order traversal approach.

To understand the need for a post/pre-order traversal method, note that, for a focal edge *e* which is alive at time *τ* before the present day, the full tree 𝒯 can be subdivided into two components: the descendants of edge *e* which we denote as the subtree 𝒯_*e*_(*τ*), and the remainder of the tree, 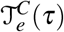, which contains all the ancestors of node *e* as well as their non-*e* descendants. For ease of writing, we call 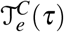 the ‘ancestral complement’ of edge *e*. Importantly, as our state model is Markovian (the type of speciation event and state-transition events depend only on the current state of the lineage), 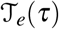 and 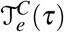 are conditionally independent given the state of edge *e* at time *τ, Y*_*e*_(*τ*). As such, we can decompose the expression for the probability *A*_*e,i*_(*τ*) as:

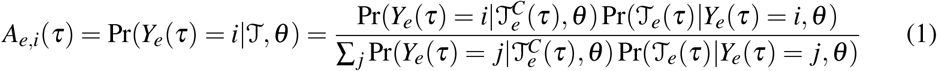

where Pr(𝒯_*e*_(*τ*) | *Y*_*e*_(*τ*) = *i, θ*) is calculated during the post-traversal step and 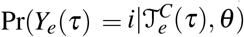 during the pre-traversal step described below (see the Supplementary Materials for a derivation of Eq. 1). Specifically, we can draw from the well-developed tree likelihoods for SSE models to calculate the probability of observing a descendant sub-tree, 𝒯 _*e*_(*τ*), given that the edge *e* is present in state *i* at time *τ* before the present day:

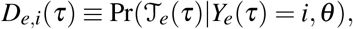

a probability that can be computed backward in time. As the probability 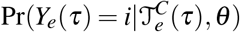 is obtained through a the pre-traversal algorithm (forward-in-time) we rewrite this expression in terms of time *t*:

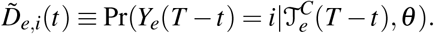

Because 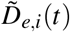 is conditioned on 𝒯^*C*^, its calculation departs from what is done in previous literature [12, 51]. However, it is again a forward-in-time probability that can be obtained using a system of ordinary differential equations. The initial conditions will depend on the backward-in-time probabilities, hence we implement the post-order algorithm followed by the pre-order algorithm. With this notation, Eq. 1 becomes

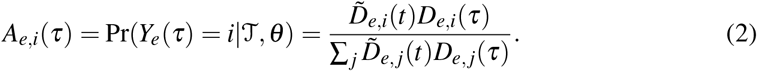

To calculate the probability that a node *N* is in state *i*, we use the time *τ*_*N*_ when the speciation event occurred, and calculate the corresponding *A*_*e,i*_(*τ*_*N*_). In summary, SAASI return a vector of *A*_*i*_ values at each node *N*, with the sum over states *i* equal to 1.

### Four-step Algorithm

#### Step 1: Post-order traversal

Here, we derive an initial value problem backward in time for *D*_*e,i*_(*τ*) –– the probability that an edge *e* in state *i* alive at time *τ* before the present day gives rise to the observed descendants between time *τ* and the present. This derivation is standard (see [12, 51, 52] for a review), but we review it here to emphasize differences with the pre-order traversal below (step 3).

We consider the change in *D*_*e,i*_(*τ*) over a small interval of time Δ*τ*, assuming that at most one event can occur.

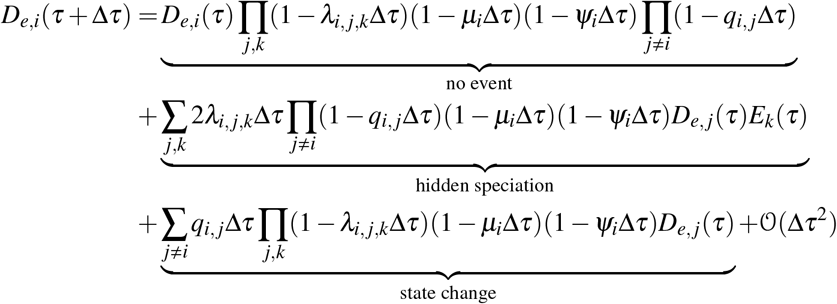

where *E*_*k*_(*τ*) is the probability that a lineage in state *k* at time *τ* has no observed descendants. Using the definition of the derivative, we obtain:

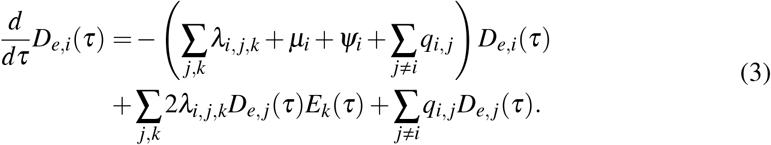

An edge *e*, which by definition does not include the node itself, can originate (at time *τ*_*e*,0_) at one of two types of nodes, either a sampling event or a speciation event where the focal edge gives rise to two descendant edges *e*_1_ and *e*_2_.

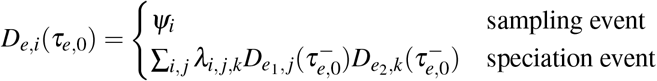

where the notation 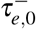 is used to emphasize that these edges are descendants (closer to the tips) of the focal edge.

To calculate *D*_*e,i*_(*τ*), we need to first solve *E*_*e,i*_(*τ*) - the probability that an edge *e* in state *i* at time *τ* has no observed descendants between time *τ* and the present day. This probability can be obtained with the following differential equation (which is derived in a similar manner as above).

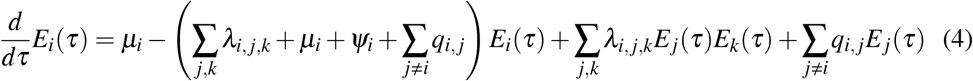

where the initial condition for *E*_*i*_(*τ*) is the probability that the focal edge is unsampled (since we are not forced to sample all the lineages or any percentage of the lineages at the present day): *E*_*i*_(0) = 1.

### Step 2: Root state probabilities

Using the initial value problem (IVP) above to solve for *D*_*e,i*_(*τ*) up the whole tree to the root, we can calculate the probability that the root is in state 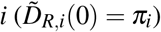 :

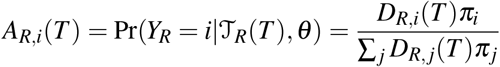

Given these root state probabilities, we can then proceed with the pre-order traversal algorithm down the tree to calculate 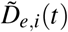.

### Step 3: Pre-order traversal

To derive an IVP for the probability 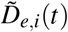, we use a similar approach as above by considering the change in this probability over a small interval of time Δ*t*. In words, we consider the probability that an edge *e* is in state *i* at time *t* + Δ*t* given the state of its immediate ancestor at time *t*. Unlike above, however, the probability 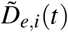 is conditioned on the observed ancestral complement tree 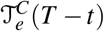. Hence along an edge *e* the only events that we should include are those that reflect a change of state (e.g., a state change or a cladogenic state change where one descendant is unobserved), not those that reflect a change of topology (since we are conditioning on the topology). In consequence, we do not include terms for birth or extinction; conditioning on the observed tree, we know that these have not occurred. This is a substantial contrast to the post-traversal step in which we computed *D*_*e,i*_(*τ*), the likelihood of 𝒯_*e*_ given that edge *e* is in state *i* at time *τ*.

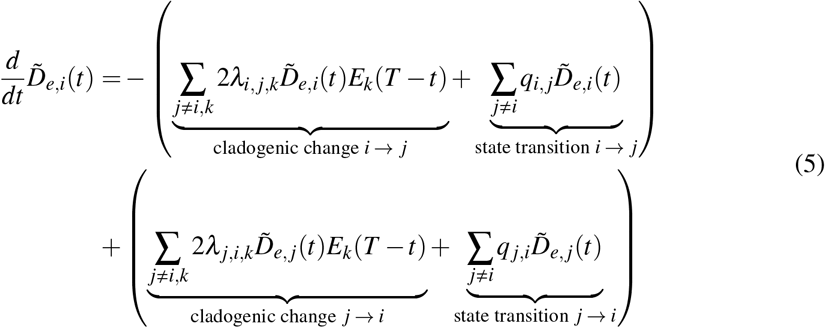

Note that the probabilities must sum to one 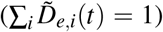 and hence this equation can be considered in terms of probability flux from one class (e.g., *i*) to another (e.g., *j*) and vice versa. The extinction probability *E*_*k*_(*T − t*) remains the same as described in Eq. 4 because the lineage will eventually go extinct and thus will not affect the tree topology.

If we consider only anagenic state change (i.e. a change of state without an accompanying speciation event), then we can further simplify the equation 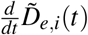 by only considering the state transition events:

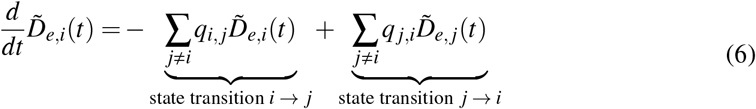

Note that this still allows for a rapid transition from one state to another immediately after a speciation event, but does not include speciation events that cause state changes. The method can readily accommodate these if the relevant *λ* rates are known. For initial conditions, note that the edge *e* can originate (forward in time at time *t*_*e*,0_) at either the root or at a speciation event. At the root, the ancestral complement 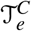 consists solely of the root itself and hence its probability is simply the prior on the root state, 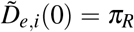. At a speciation event, the state of the focal edge *e* at time *t*^+^ (i.e., immediately following the speciation event) depends on (1) the state of the parent lineage, edge *e*^*−*^, immediately prior to the speciation event at time *t*^*−*^, (2) the type of speciation event to occur as given by the probability 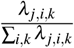, and (3) the state of the sister edge *e*^*′*^ of the focal edge immediately following the speciation event. Specifically,

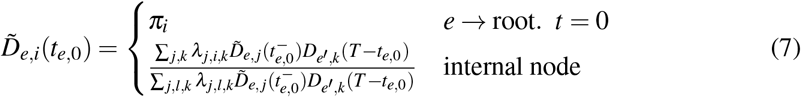

Note that in Eq. 7, the backward-in-time probabilities *D*_*e*_^*′*^_,*k*_(*T − t*_*e*,0_) for the sister edge *e*^*′*^ are evaluated at the end of the branch. The necessary values were previously calculated from time *t*_*e*,0_ during the post-order traversal and sorted for use in initializing the pre-order traversal at internal nodes.

We therefore solve Eq. 5 along each edge, forward in time. Upon reaching an internal node *N* of the phylogeny, we use Eq. 7 to initialize the ODE on the two edges (say *e*_1_ and *e*_2_) descending from *n*. At this point, note that the ancestral complement tree, on which we are conditioning, changes from 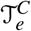 to 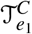 or 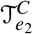. Both of these include node *N*.

### Step 4: Ancestral State Probabilities

Once 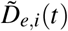 is computed during the pre-traversal step the ancestral state probabilities can be computed via the product outlined above:

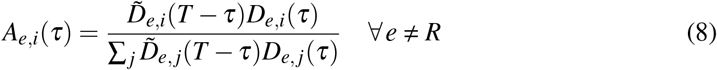

At each internal node *N* we then define the “reconstructed state” as the state with the highest probabilities at those nodes:

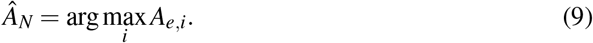

### Parameter estimation

Since SAASI requires the users to know the speciation, extinction, sampling and transition rates as their inputs, these rates need to be estimated. We estimate the speciation and extinction rates using the maximum likelihood method proposed by [53], where we assume that the sampling rate is known. To estimate the transition rates in the simulations, we use ‘ER’ model implemented in ace. The ‘ER’ model assumes that all the transition rates are equal, which is one of the parameter settings in the simulations (and not a requirement of SAASI). We note that in principle we could estimate input rates with BiSSE, but there is no readily available implementation allowing for non-ultrametric trees, and [54] reported some limits to BiSSE estimation for larger phylogenies.

We compared the ‘ER’ model’s transition rates to the truth using simulated trees (see below), and found that ace infers the transition rates accurately if there is little to no sampling difference between states. However, if state *i* is far less sampled than state *j*, ace overestimates the transition rates from other states *j* to state *i* (*q* _*ji*_) and underestimates transition rates from state *i* to other states *j* (*q*_*ij*_). The error depends on the sampling ratios. Therefore, we also test how transition rate adjustments affect our inferences, scaling *q*_*ij*_ and *q* _*ji*_ based on our simulated findings.

For state *i*, we simultaneously adjust 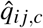 and 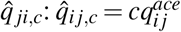 and 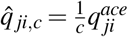, where *c* is the multiplicative factor that adjusts the transition rates and 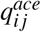 is the transition rate estimated from ‘ace’. We refer to this transition rate adjustment as “adjusting state *i* by a factor of *c*”.

### Simulation tests

#### Single demonstration tree

We first use an illustrative example in which we compare ancestral state inference using the SAASI versus ace on a simulated tree with binary states. Specifically, we consider a case in which the states differ only in their sampling rates, with state 1 being sampled at one-tenth the rate of state 2. All other diversification rates are identical between the states (*λ*_1_ = *λ*_2_ = 1, *µ*_1_ = *µ*_2_ = 0.045, *q*_12_ = *q*_21_ = 0.05 per unit time).

We assume that the root is in state 1, reflecting, for example, a scenario where the place of origin of an outbreak has a lower sampling rate than other locations. The resulting tree has 87 tips, of which only 9 (10.3%) are in state 1 (Figure 1, a). We perform SAASI using the true parameters as inputs and ace using the equal rate (‘ER’) model. We further examine how our accuracy would change if the sampling rates are mis-specified, by varying the sampling rates *ψ*_1_ and *ψ*_2_ (see Supplementary Fig. 3). We consider different sampling ratios 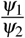, where the ratio ranges from 0.1 (the true sampling rates) to 2.0 (state 2 is incorrectly assumed to be sampled at one-half the rate of state 1).

#### 1,000 Simulated trees under different sampling scenarios

To evaluate SAASI’s performance, we simulated 1,000 trees with two states under five scenarios with varying intensities of sampling and bias correction strategies:

1. High sampling bias: state 1 is sampled at one-tenth the rate of state 2 (*ψ*_1_ = 0.1 and *ψ*_2_ = 1.0).
2. Moderate sampling bias: state 1 is sampled one-fourth the rate of state 2 (*ψ*_1_ = 0.25 and *ψ*_2_ = 1.
3. High sampling bias with downsampling: after generating trees with *ψ*_2_ = 10*ψ*_1_, we randomly downsample tips from state 2 to achieve uniform sampling (preserving 10% of the state 2 tips to match state 1’s sample size).
4. Moderate sampling bias with downsampling: after generating trees with *ψ*_2_ = 4*ψ*_1_, we randomly downsample tips from state 2 to achieve uniform sampling (preserving 25% of the state 2 tips to match state 1’s sample size).
5. Uniform sampling: state 1 is sampled at the same rate as state 2 (*ψ*_1_ = *ψ*_2_ = 0.5).

Trees are generated with state-dependent speciation rates *λ*_1_ = 3 and *λ*_2_ = 1.5, extinction rates *µ*_1_ = 0.1 and *µ*_2_ = 0.05, and transition rates *q*_12_ = *q*_21_ = 0.3. We generate trees with a maximum running time of 3,000 units and a maximum of 3,000 tips.

For scenarios (1), (2), and (5), we perform ancestral state inference using ace (equal rates model) [22], PastML (MPPA method with F81 evolutionary model) [26], simmap (equal rates model, but set the equilibrium state frequency to match the sampling ratios) [20], TreeTime (mugration model) [27], TreeTime (mugration model with sampling bias correction), SAASI with true parameters, and SAASI with estimated parameters (SAASI^*∗*^). In SAASI^*∗*^, we assume the sampling rates are known, speciation and extinction rates are estimated based on [53], and transition rates are estimated using ace. For the downsampling scenarios (3) and (4), we apply only ace, PastML, simmap, and TreeTime without sampling correction, as these methods assume uniform sampling. For each simulation, we recorded the wall-clock time for each method.

#### Scalability

To evaluate SAASI’s computational performance across different inference methods, we record the computation time for each method from the previous scenarios (1,000 simulated trees under different sampling scenarios). To evaluate SAASI’s computational performance across different tree sizes and number of states, we simulate trees with two states ranging from 100 tips to 100,000 tips using the following parameter values: *λ*_1_ = 3, *λ*_2_ = 1.5, *µ*_1_ = *µ*_2_ = 0.045, *q*_12_ = *q*_21_ = 0.3, *ψ*_1_ = 0.1 and *ψ*_2_ = 1.0. This results in tree sizes of 100, 200, 500, 1,000, 2,000, 5,000, 10,000, 50,000, and 100,000 tips. We also evaluate SAASI’s computational performance across different numbers of states *k* between *k* = 2 and *k* = 30. For each state, we set the following parameter values: *λ*_*i*_ = 3, *µ*_*i*_ = 0.05, *q*_*ij*_ = 0.3, *ψ*_*i*_ = 0.1. We record the wall-clock time required for each method to complete ancestral state inference.

### Accuracy comparison

We consider two measures of accuracy: “consensus accuracy” *a*_consensus_ and “probability accuracy” *a*_probability_. Consensus accuracy is the fraction of internal states that are inferred correctly, when each node’s inferred state is its highest-probability state (*i* for which *A*_*e,i*_(*τ*) in Eq. 8 is maximized). In contrast, probability accuracy weights each node by the inferred probabilities of the true ancestral state: 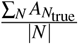, where 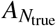 is the inferred probability of the true state at node *N* and we sum over the number of internal nodes. For example, if a tree had only one internal node and its true state was ‘state 1’, and Eq. 8 gave state 1 a probability of 0.51, the consensus accuracy for that tree would be 1, and the probability accuracy would be 0.51.

For downsampling scenarios (3) and (4) from B, we calculate accuracy in three ways to evaluate the trade-offs of downsampling:

1. Downsampled only: accuracy calculated only on internal nodes present in the downsampled tree. This represents the best-case scenario for downsampling, where only preserved nodes are evaluated, and no attention is paid to the loss resulting from removing data.
2. Downsampled with missing transitions: accuracy calculated on all internal nodes from the original pre-downsampling tree, but counting nodes that are removed during downsampling as not having accurate inferences. This represents the cost of downsampling in terms of accuracy loss.
3. No downsampling: accuracy calculated on the complete original tree using all seven methods (ace, PastML, simmap, TreeTime, TreeTime with correction, SAASI, SAASI^*∗*^). This serves as a baseline comparison to evaluate whether downsampling or sampling correction methods yield better results.

### Application to avian influenza H5N1

We apply SAASI to a timed phylogeny from the highly pathogenic outbreak of avian influenza A (H5N1; hemagglutinin (HA) segment) in US dairy cattle [2]. The sample collection dates ranged from April 2023 to April 2024, with 104 sequences. We use this phylogeny to illustrate how accounting for sampling differences between wild birds and other species can affect inferences about the initial spillover event and the subsequent transmissions among host species. Since the influenza virus spread both geographically and across multiple host species, we use SAASI to compare reconstructions of host movements and geographical movements under several plausible sampling adjustments.

This particular phylogeny (Supplementary Fig. 2) in [2]; `mcc-gtrg-ucld-gmrf-colored.pruned.tre`) is a portion of a Bayesian time-calibrated phylogeny. It contains sequences from 6 taxonomic groups (cattle, (wild) mammal, domestic mammal, poultry, human, and wild bird) and 12 sampling locations (California, Indiana, Kansas, Maryland, Michigan, Minnesota, Montana, New Mexico, North Carolina, Ohio, Oklahoma, Texas and Wisconsin). We excluded the single human sample from our analysis. We also combined mammal and domestic mammal samples into one group.

In their study [2], the authors inferred that the H5N1 cattle outbreak was likely due to a single spillover event from wild birds to cattle, followed by transmission within the cattle and subsequent transmission between species. They estimated that the first spillover event occurred on December 9, 2023, with 95% highest posterior density (HPD) between October 12, 2023 and January 26, 2024. Their spatial and host phylogeographic analysis was conducted using BEAST using BSSVS [5, 41].

We estimate the time of the first spillover event from the wild bird population to cattle and assess the robustness of the time estimate to the relative sampling. We note that in our context, “sampling” refers to the fraction of infections that are represented in the data (as opposed to the fraction of the population that is sampled, or the number of samples). Accordingly, wild bird infections (wild birds being the normal host population for avian influenza viruses) may have a much lower sampling rate than, for example, infections in domestic cattle.

#### H5N1 host analysis

We consider three scenarios to capture different levels of sampling bias. First, we model no significant difference in sampling between wild birds and other species. Although this scenario is unlikely, it serves as a natural baseline and reflects the assumption commonly made in phylogeographic methods such as ace or simmap. In the second scenario, we introduce a moderate sampling bias, modelling wild bird infections as being sampled ten times less frequently than H5N1 infections in other species. Finally, the third scenario reflects a severe sampling bias, in which wild bird infections are sampled one hundred times less frequently than infections in other species. Wild bird populations are challenging to sample, and the probability of sampling any given H5N1 infection (compared to that in cattle) is unknown. This motivates our wide range of sampling differences.

We then explore the inferred ancestral state of the clade of interest and reconstructed interspecific transmission events under different sampling rates using SAASI, which requires us to estimate the speciation, extinction, and transition rates. We adapted the estimated transition rates from [55]. The mean transition rate between non-wild bird hosts was two transitions per year, while the transition rate from wild birds to other hosts was higher, with a mean of 4 transitions per year. We further assume that the transition rate from wild birds to the other hosts is two times higher (than this mean) and that the transition rate from other hosts to wild birds is two times lower than this mean, informed by our simulation results for how sampling affects estimated transition rates. Hence, we obtain the following transition rate matrix:

|  | Cattle | Mammal | Poultry | Wild bird |
| --- | --- | --- | --- | --- |
| Cattle | — | 2 | 2 | 1 |
| Mammal | 2 | — | 2 | 1 |
| Poultry | 2 | 2 | — | 1 |
| Wild bird | 4 | 4 | 4 | — |

These transition rates are similar to those estimated using the ace symmetric transition rate, which has a mean of 3.68 per year. We assume that the speciation and extinction rates for all species are the same, with estimated rates of 21.1 and 6.8 per year, respectively.

To estimate the speciation and extinction rates, we used the method proposed by [53], which calculates the likelihood of the tree. The method assumes the tree is generated by the birth-death-sampling process and uses bifurcation time and sampling time. This method does not consider state-dependent events. Thus, our speciation and extinction rates only have one value. We set the speciation and extinction rates equal for all the states to show the effect of sampling rate differences.

#### H5N1 geographic analysis

We use geographic locations to explore the impact of different geographic sampling rates on inferences about the likely geographic origin of the outbreak, focusing on the geographic location of the transmission event(s) from wild birds to cattle. Given that most of the samples were collected in Texas, we consider two alternative sampling models in the phylogeographic analysis. In the first, baseline model, we assume no significant difference in sampling effort between Texas and other states; that is, all states are modelled as having the same sampling rate. In the second model, we account for sampling bias by assuming that Texas was sampled five times more frequently than other states. Although we do not know the true relative sampling differences between the locations, this analysis will illustrate how sensitive the inferred geographic location of the spillover event is to sampling differences.

In this analysis, because the transition rates are unknown and difficult to estimate, we model that the transition rates are equal between states, and estimate them using ace with the ‘ER’ model. We use the the speciation and extinction rates estimated by the species analysis.

## Acknowledgements

We emphasize our strong appreciation for the authors of Nguyen et al. [2], the Flu crew at the U.S. National Animal Disease Center and the authors of the GitHub repository at https://github.com/flu-crew/dairy-cattle-hpai-2024, who published their H5N1 repository under a license permitting reuse and publication without restriction (among other permissions). Such data sharing is essential for method development efforts, and it is greatly appreciated.

## Author contributions

Y.S. contributed conception or design of the work; analysis; interpretation of data; drafting and revision of the manuscript. I.G. contributed the creation of new software used in the work and revision of the manuscript. A.M. contributed conception or design of the work; analysis; drafting and revision of the manuscript. C.C. contributed conception or design of the work; analysis; drafting and revision of the manuscript.

## Data Availability

Data that support the findings of this study can be found in [2]: https://github.com/flu-crew/dairy-cattle-hpai-2024/tree/main: mcc-gtrg-ucld-gmrf-colored.pruned.tre.

## Code Availability

Code that support the findings of this study have been deposited in: https://github.com/yexuansong/saasi_manuscript. Source Data are provided with this paper. SAASI is available as an R package at https://github.com/MAGPIE-SFU/saasi. Given the difficulty of specifying a full *λ*_*ijk*_ matrix, the current implementation is as in Eq. 6 (*λ*_*ijk*_ = 0 if *i* : *j, k*).

## Sampling Aware Ancestral State Inference

### Supplementary Information

#### Derivation of the main decomposition

For clarity, we provide an explicit derivation of our central equation for the *A*_*e,i*_,

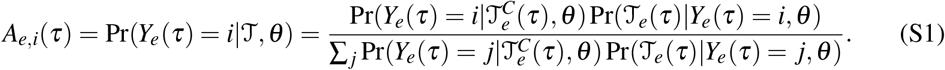

In this derivation, we will suppress *τ* and *θ* to simplify notation. For example, we write 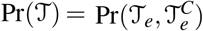, and *A*_*e,i*_ = Pr(*Y*_*e*_ = *i*|𝒯). By Bayes’ theorem, we have

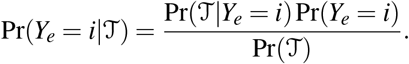

Since 𝒯_*e*_ and 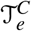 are conditionally independent given *i* (which means, given that edge *e* is in state *i* at time *τ*), the above is

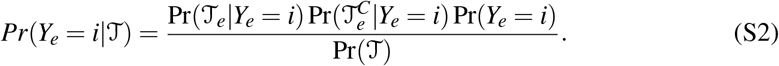

Using Bayes’ theorem again, we write

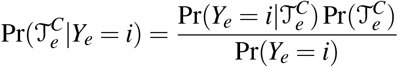

and

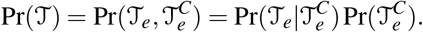

Substituting these into Eq. S2, we have

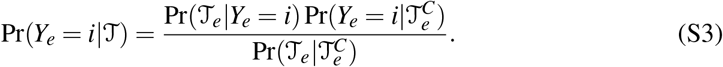

It remains to show that the denominator has the form given in (1) of the main text. Since the state of edge *e* must be one and only one of the possible ancestral states, we have

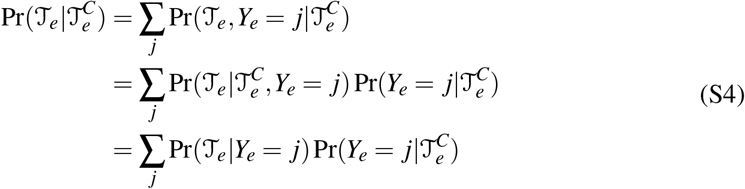

where the second line is a conditional probability, and the last line is due to the conditional independence of T_*e*_ and 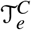 given the state of edge *e*.

## Supplementary Figures

**Supplementary Fig. 1** gives an example illustrating that SAASI’s result is very similar to ace’s if we assume equal sampling rates. In other words, while SAASI’s underlying mathematical model is quite different from that of ace, the results are very similar if the same assumption is made about sampling. The natural comparison for this figure is Figure 1 in the main text.

**Supplementary Fig. 2** shows how the consensus and probability accuracy changes if the sampling ratio is misspecified. The accuracy is sensitive to the sampling rates. Both accuracies remain above 0.75 for sampling ratios *ψ*_1_/*ψ*_2_ *∈* [0.15, 1]. If the sampling rate is strongly mis-specified (with *ψ*_1_/*ψ*_2_ > 1 reflecting the assumption that state 1 is *more* highly sampled than state 2, rather than less), then the accuracies decrease to a level similar to the accuracy from ace, i.e. just over 0.55.

**Supplementary Fig. 3** compares the accuracies obtained using different ancestral state inference methods and adjustments to the transition rates, including mis-specification of the transition rates. SAASI with true rates has the highest accuracy, and SAASI’s results under various transition rate adjustments are similar to one another. This demonstrates the robustness of SAASI to misspecification of the transition rate.

**Supplementary Fig. 4** compares ancestral state inference using ace and SAASI under the assumption of equal sampling in all hosts. Neither method can identify the key transition events from wild birds to cattle in this circumstance, nor reliably infer the internal node states before November 2023.

**Supplementary Fig. 5** compares ancestral state inference using SAASI under the assumption that wild bird infections are sampled 100 times less than infections in other host species, with equal transition rates. The results are similar to those in the main text, where we adjusted the transition rates to and from wild birds to other species by a factor of 2.

**Supplementary Fig. 6** compares ancestral state inference methods under moderately biased sampling. All methods have high consensus and probability accuracies; SAASI and SAASI^*∗*^ have smaller tails.

**Supplementary Fig. 7** compares ancestral state inference methods under highly biased sampling in different scenarios. If the tree is downsampled under the specified sampling ratio, then both the consensus and probability accuracy are high, considering only the sampled nodes. However, if the missing transitions are taken into account, then the accuracies decrease due to missing data.

**Supplementary Fig. 8** compares ancestral state inference methods under moderately biased sampling in different scenarios. Similarly to Supplementary Fig. 6, if the tree is downsampled under the specified sampling ratio, then both the consensus and probability accuracy are high, considering only the sampled nodes. However, if the missing transitions are taken into account, then the accuracies decrease due to missing data.

**Supplementary Fig. 9** compares ancestral state inference methods under uniform sampling. All methods achieve high accuracy.

**Supplementary Fig. 10** compares the running time of ancestral state inference methods. ace, PastML, and TreeTime have fast running times and can perform inference for a tree of size 1,600 in less than 10 seconds. On the other hand, simmap, SAASI, and SAASI^*∗*^ have running times longer than Markov-based methods, with completion times ranging from 20 to 35 seconds. The running time increases with the size of the tree.

**Supplementary Fig. 11** compares SAASI’s running time for different tree sizes (with two states). The running time grows linearly with the number of taxa. Increasing the number of states also increases the run time.

**Supplementary Fig. 12** compares SAASI’s running time for different number of states *k* (from *k* = 2 to *k* = 30).

---

**Supplementary Fig. 1.**
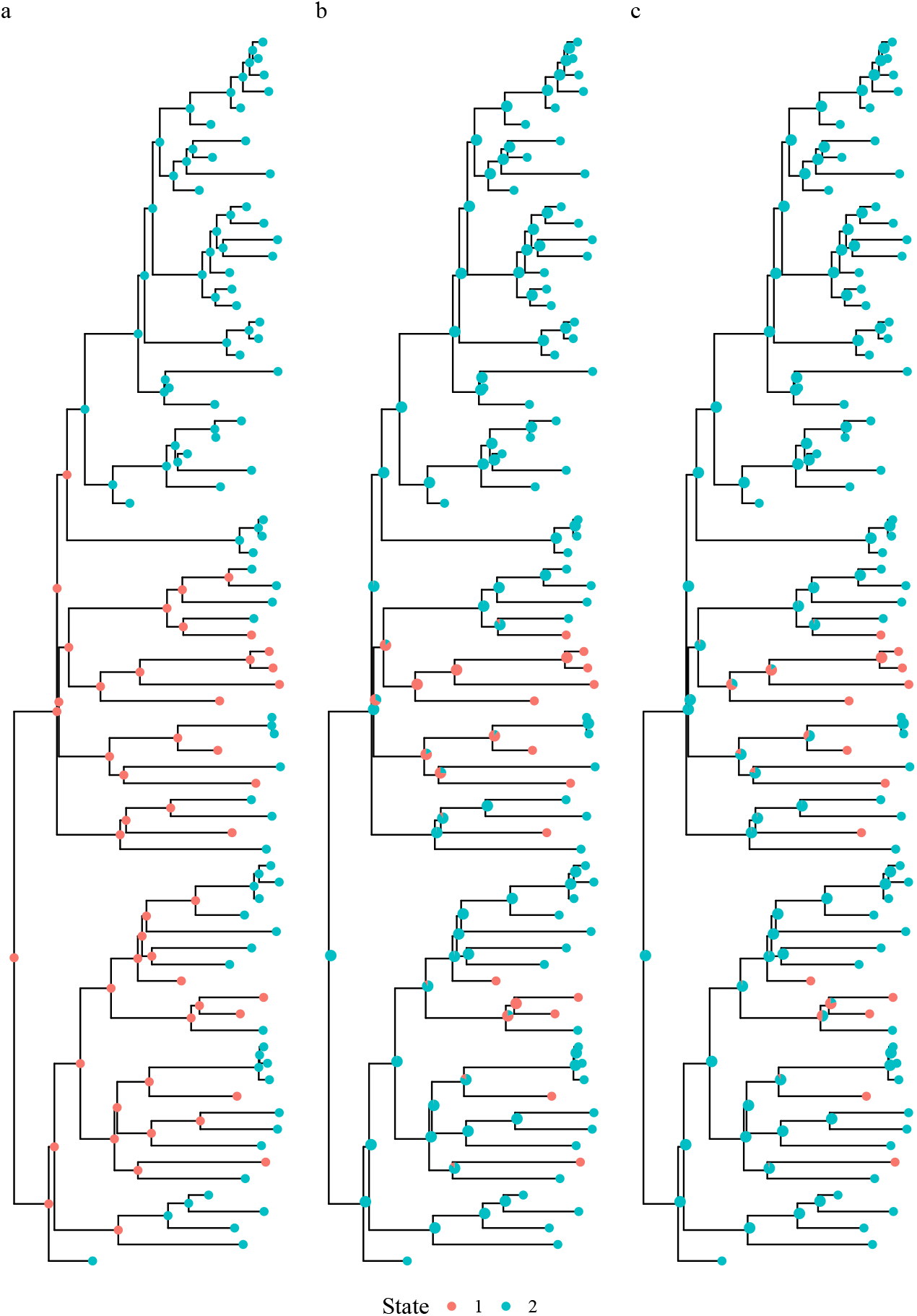
Ancestral state inference using SAASI under an equal sampling model and using ace. a: Simulated tree with known transmission histories; b: SAASI with equal sampling rates (*ψ*_1_ = *ψ*_2_ = 0.5); c: ace under equal sampling rate model. The tree is generated using the following parameters: *λ*_1_ = *λ*_2_ = 1, *µ*_1_ = *µ*_2_ = 0.045, *q*_12_ = *q*_21_ = 0.05, and *ψ*_1_ = 0.05, *ψ*_2_ = 0.5.

**Supplementary Fig. 2.**
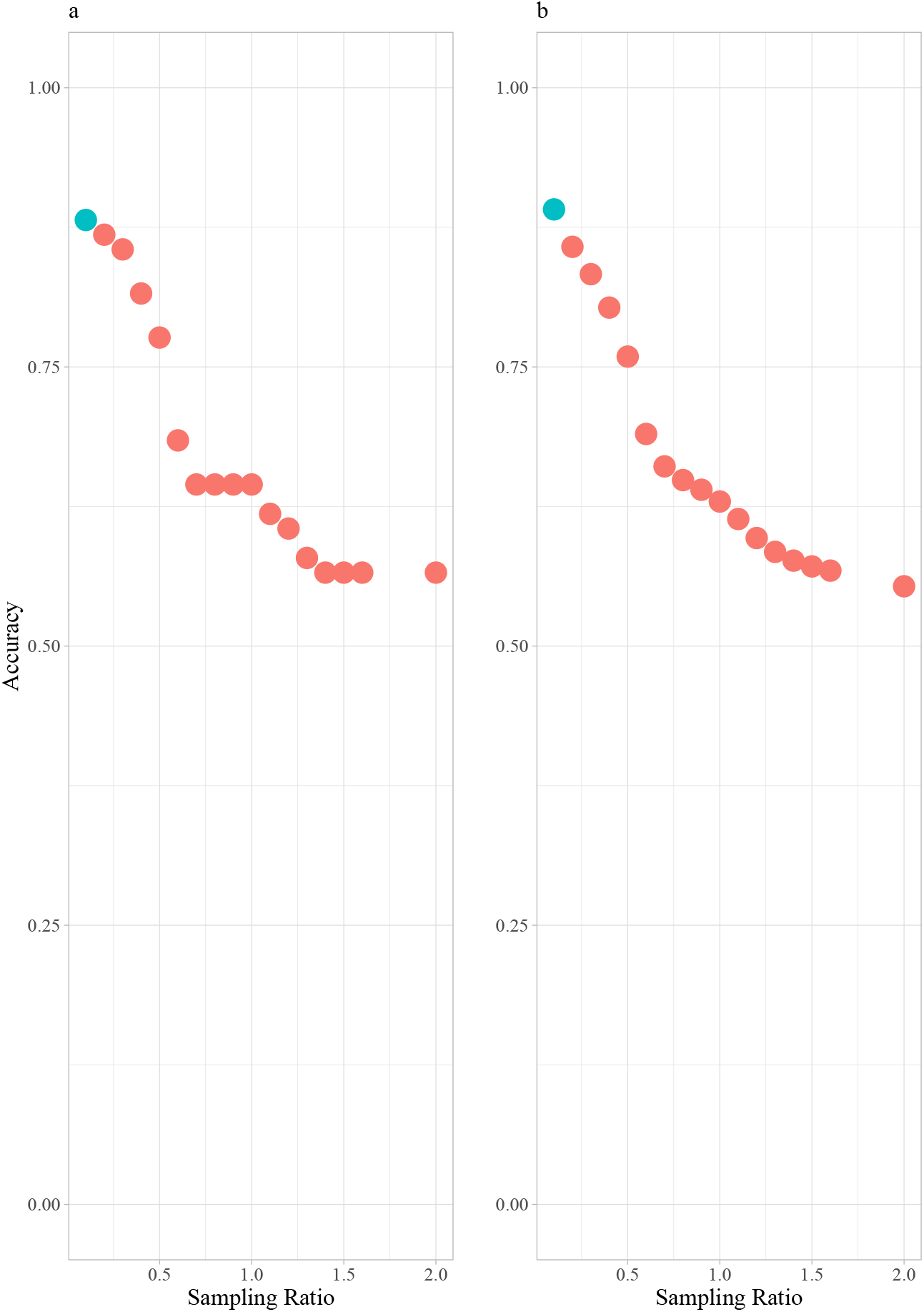
Accuracies of SAASI under varying sampling ratios on a fixed simulated tree. a: Consensus accuracy; b: Probability accuracy. The blue point represents accuracy under the true sampling ratio 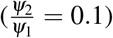. Red points represent accuracy under mis-specified sampling ratios ranging from 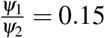.

**Supplementary Fig. 3.**
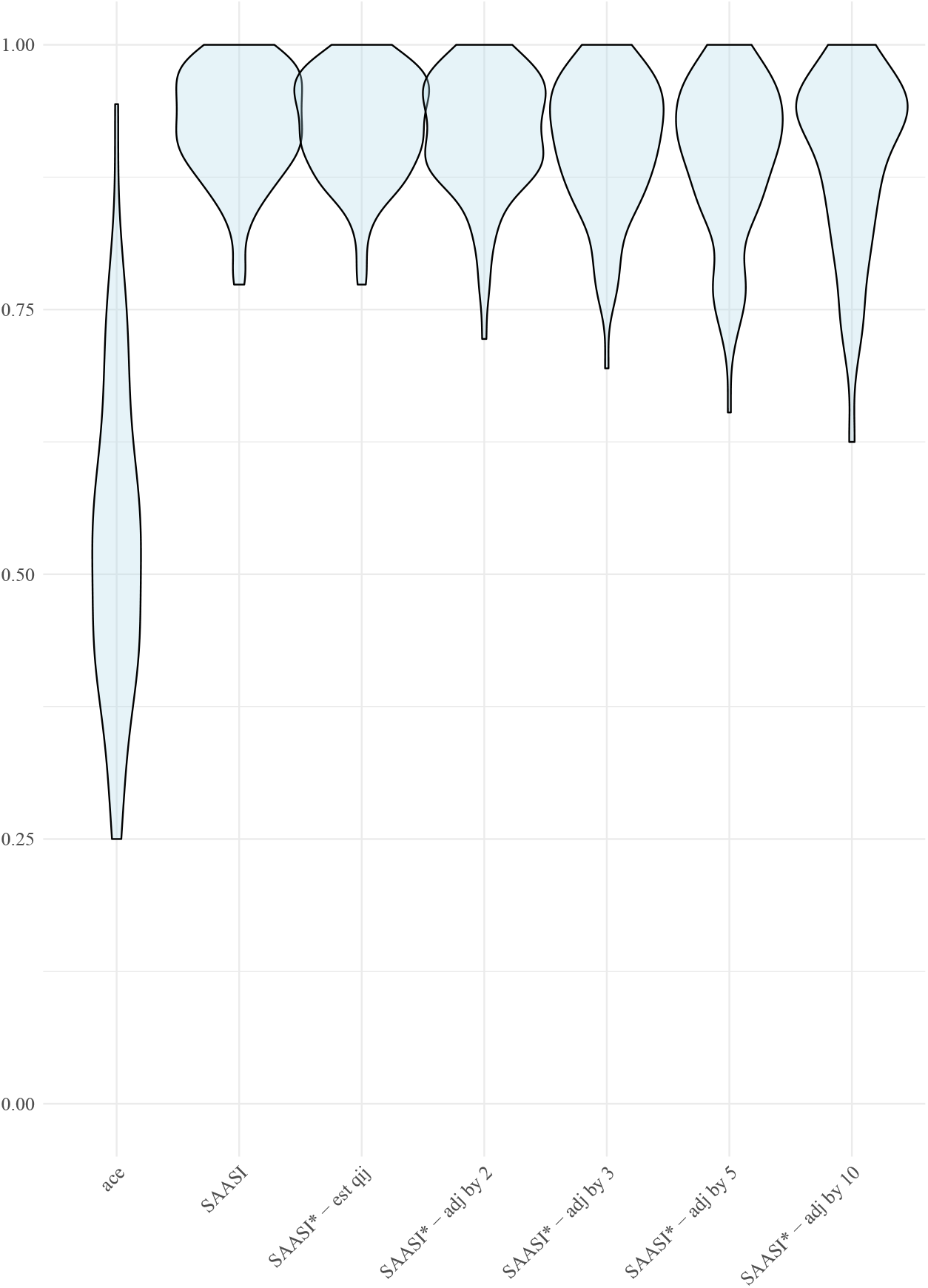
Comparison of consensus accuracies for ancestral state inference using ace and SAASI under various transition rates adjustments. state 1 is sampled 10 times less than the other states (three states in total). Simulations use *q*_*ij*_ = 0.2, *∀i, j*. From left to right: ace; SAASI with true rates; SAASI^*∗*^ with estimated transition rates from ‘ace’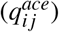; SAASI with adjusted transition rates of state 1 by a factor of *c*, ranging from *c* = {2, 3, 5, 10}.

**Supplementary Fig. 4.**
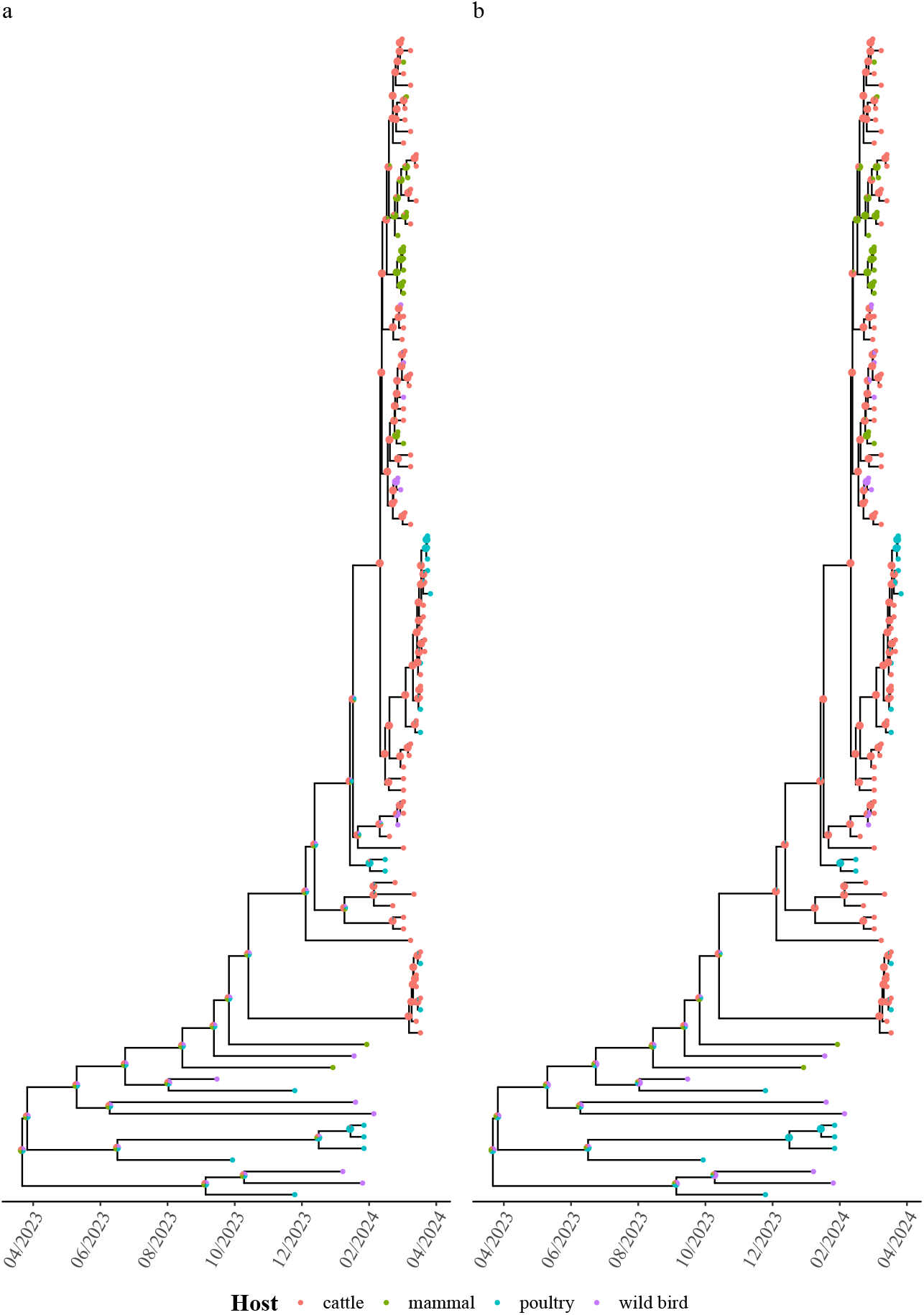
Ancestral state inference of the H5N1 HA segment tree using ace and SAASI, assuming equal sampling across species. a: ace; b: SAASI, equal sampling. Pie charts indicate the inferred probabilities of being in particular states.

**Supplementary Fig. 5.**
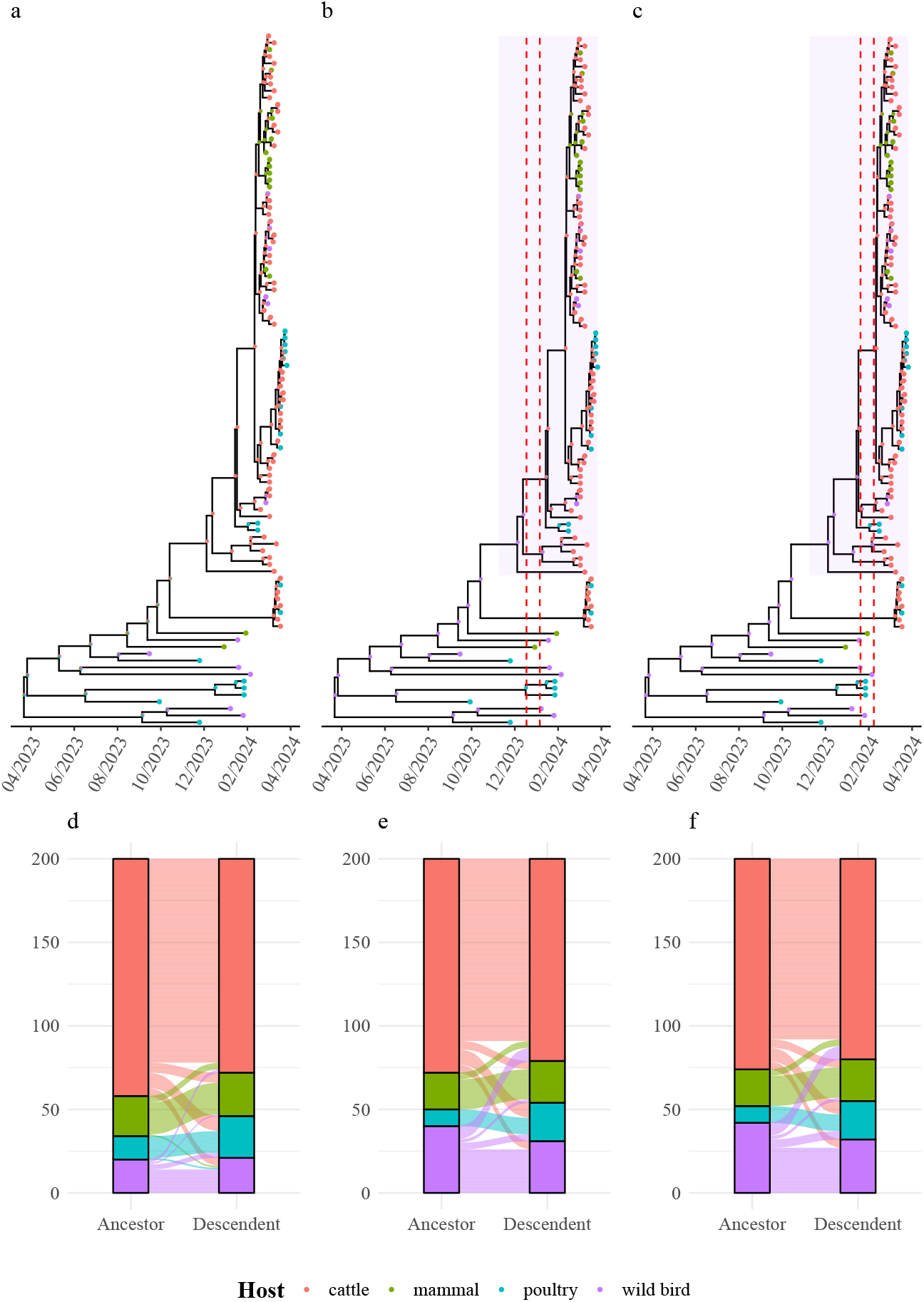
Ancestral state inference of the H5N1 HA segment tree using SAASI under different species-level sampling models. a: Inferred species hosts under equal sampling rates; b: Wild birds at one-tenth sampling; c: Wild birds at one-one hundredth sampling; d: Inferred viral transitions between host species in a; e: Inferred viral transitions between host species in b; f: Inferred viral transitions between host species in c. Pie charts indicate the inferred probabilities of being in particular states. Transition rates are equal between species. The dashed red lines indicate the key transition event from wild bird to cattle.

**Supplementary Fig. 6.**
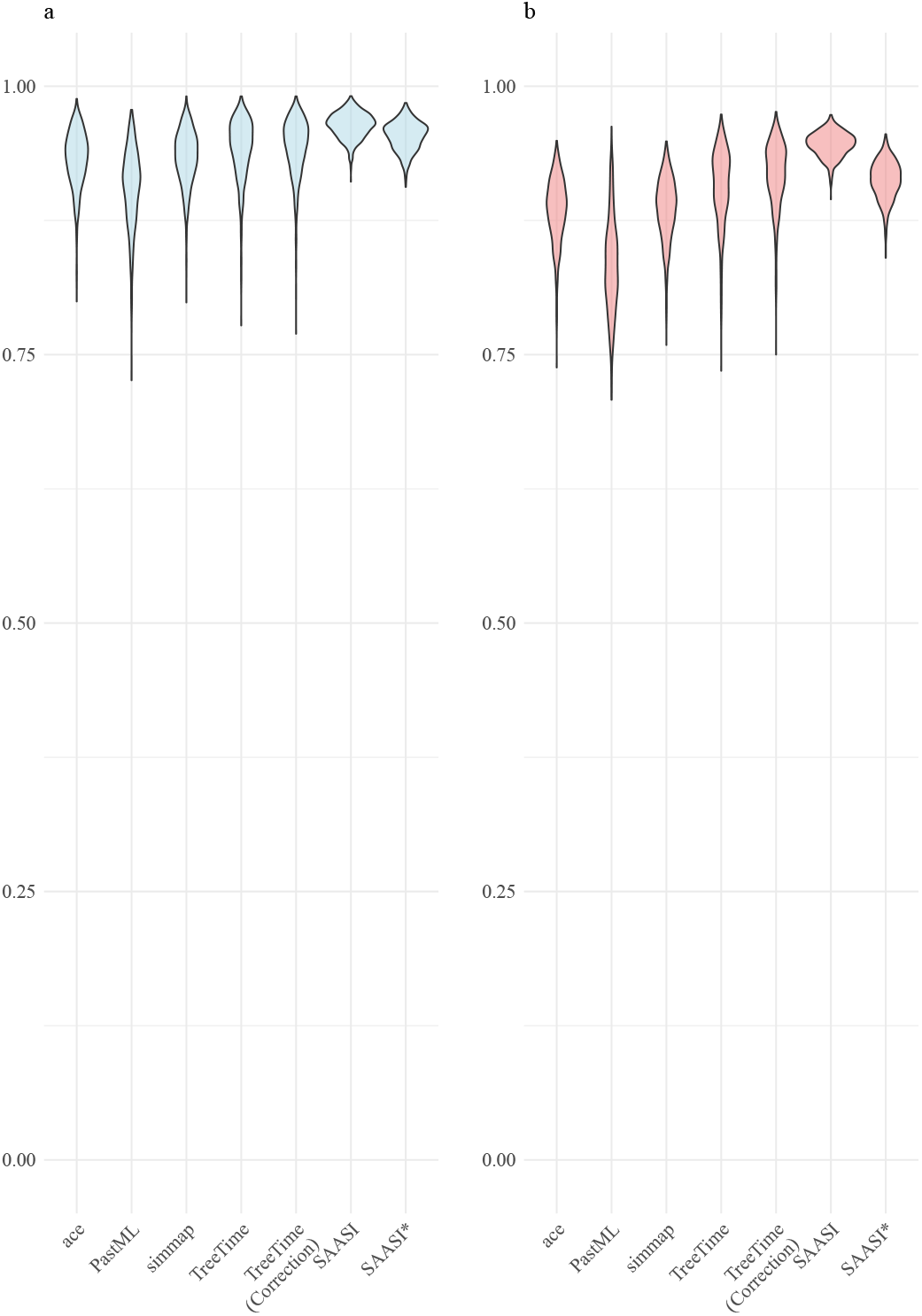
Accuracy of ancestral state reconstruction methods under moderately biased sampling (*ψ*_2_ = 4*ψ*_1_) and state dependent diversification across 1000 simulated trees. a: Consensus accuracy, defined as the fraction of correctly inferred ancestral node states. b: Probability accuracy, accounting for uncertainty in the node inference. Violin plots show the distribution of accuracy values across all simulations. ace and simmap use the equal-rates (ER) transition model. PastML employs the MPPA method with the F81 evolutionary model. TreeTime uses the mugration model with and without sampling bias correction (shown in parentheses). SAASI uses the true parameter values that generated the tree. SAASI^*∗*^ uses estimated speciation and extinction parameters but is provided with the true sampling rate. Trees were generated with speciation rates *λ*_1_ = 3 and *λ*_2_ = 1.5, extinction rates *µ*_1_ = 0.1 and *µ*_2_ = 0.05, transition rates *q*_12_ = *q*_21_ = 0.3, and sampling rates *ψ*_1_ = 0.25 and *ψ*_2_ = 1.0.

**Supplementary Fig. 7.**
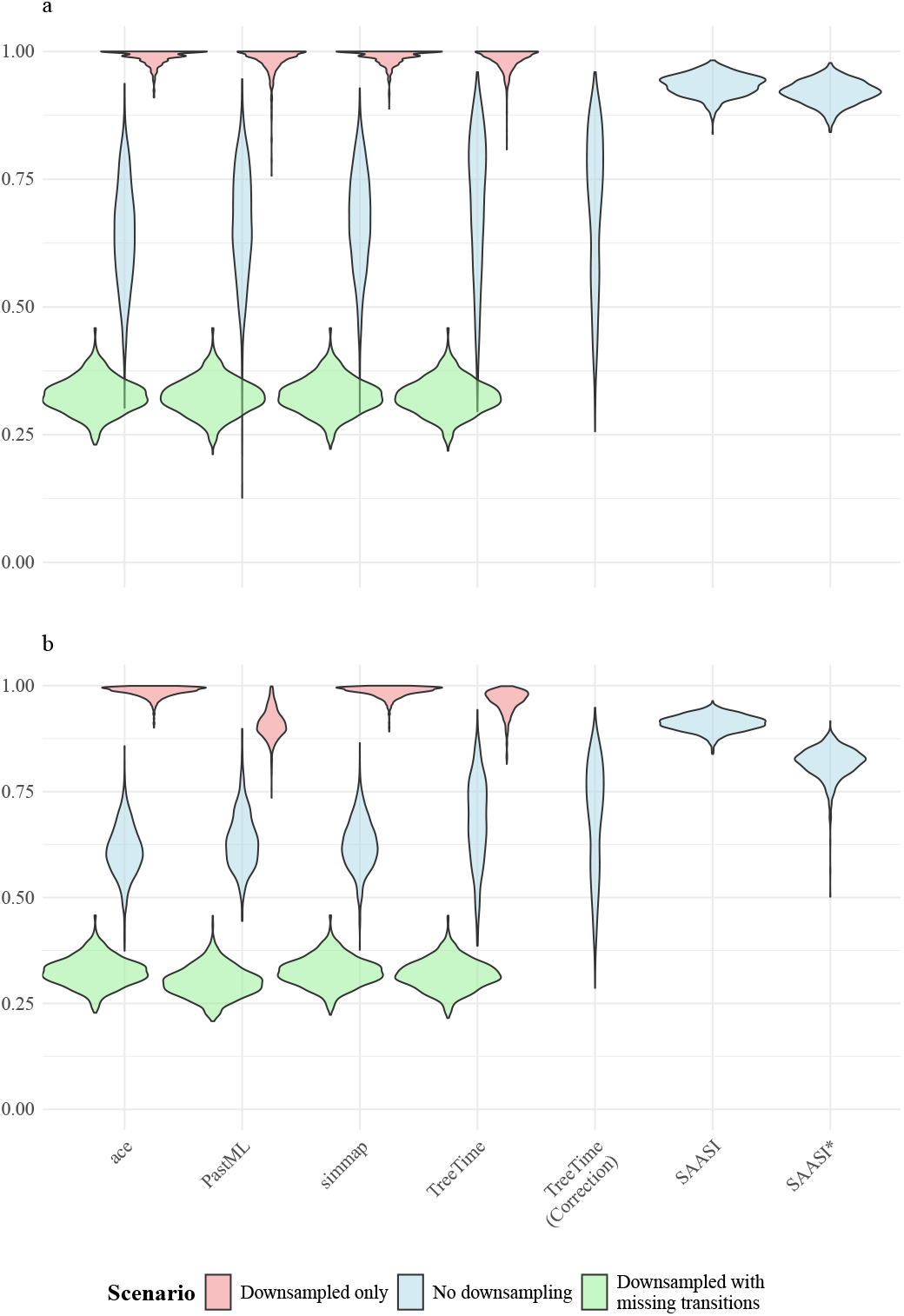
Impact of downsampling on ancestral state reconstruction accuracy under highly biased sampling (*ψ*_2_ = 10*ψ*_1_) and state dependent diversification across 1000 simulated trees. a: Consensus accuracy, defined as the fraction of correctly inferred ancestral node states. b: Probability accuracy, accounting for uncertainty in the node inference. Violin plots show the distribution of accuracy values across all simulations for three scenarios: (1) Downsampled only (red): accuracy calculated only on nodes present in the downsampled tree; (2) No downsampling (blue): accuracy on the original complete tree without any downsampling; (3) Downsampled with missing transitions (green): accuracy includes both downsampled nodes and transition events that occurred on branches removed during downsampling;. ace and simmap use the equal-rates (ER) transition model. PastML employs the MPPA method with the F81 evolutionary model. TreeTime uses the mugration model with and without sampling bias correction (shown in parentheses). SAASI uses the true parameter values that generated the tree. SAASI^*∗*^ uses estimated speciation and extinction parameters but is provided with the true sampling rate. Trees were generated with speciation rates *λ*_1_ = 3 and *λ*_2_ = 1.5, extinction rates *µ*_1_ = 0.05 and *µ*_2_ = 0.1, transition rates *q*_12_ = *q*_21_ = 0.3, and sampling rates *ψ*_1_ = 0.1 and *ψ*_2_ = 1.0.

**Supplementary Fig. 8.**
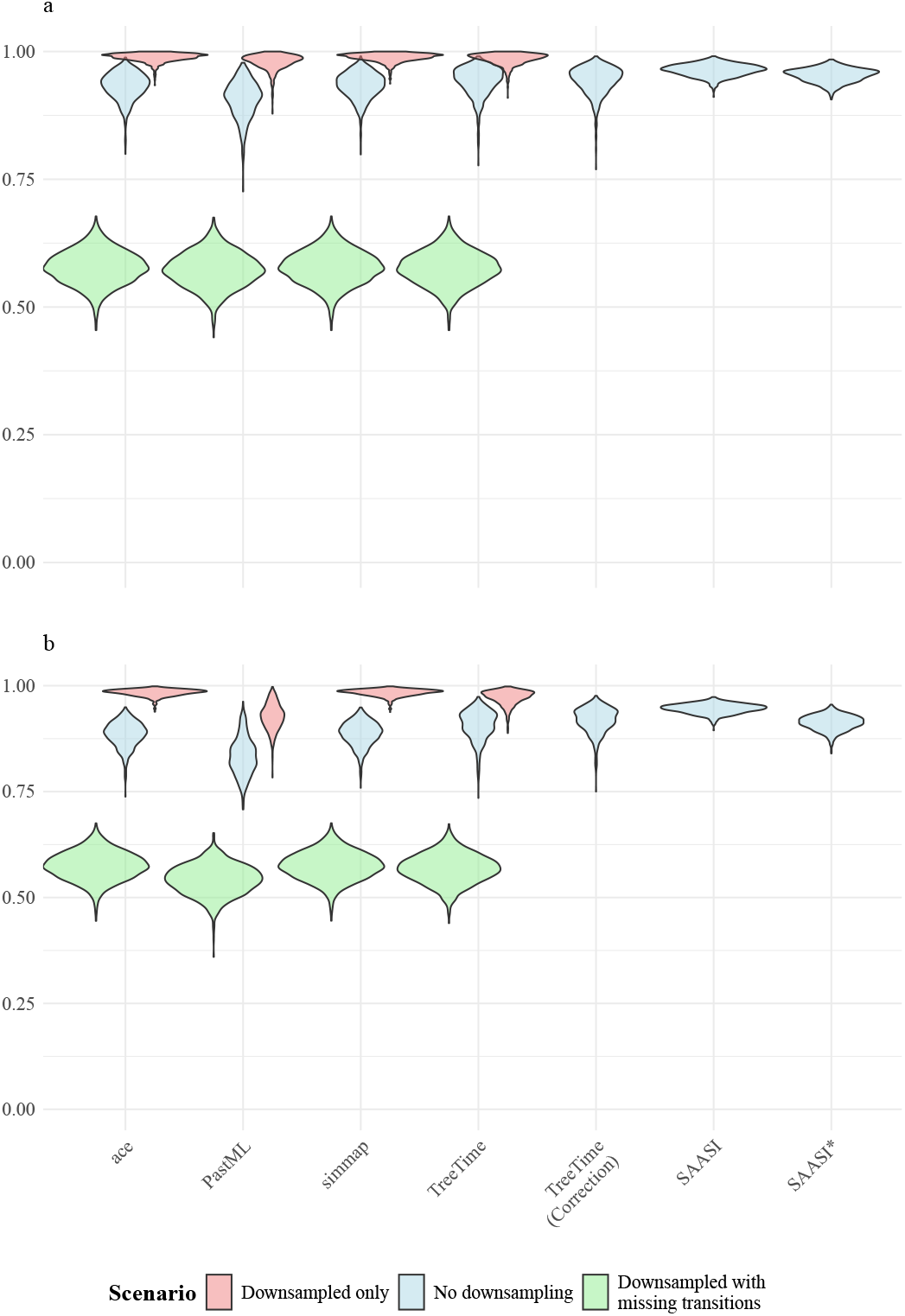
Impact of downsampling on ancestral state reconstruction accuracy under moderate biased sampling (*ψ*_2_ = 4*ψ*_1_) and state dependent diversification across 1000 simulated trees. a: Consensus accuracy, defined as the fraction of correctly inferred ancestral node states. b: Probability accuracy, accounting for uncertainty in the node inference. Violin plots show the distribution of accuracy values across all simulations for three scenarios: (1) Downsampled only (red): accuracy calculated only on nodes present in the down-sampled tree; (2) Downsampled with missing transitions (green): accuracy includes both down-sampled nodes and transition events that occurred on branches removed during downsampling; (3) No downsampling (blue): accuracy on the original complete tree without any downsampling. ace and simmap use the equal-rates (ER) transition model. PastML employs the MPPA method with the F81 evolutionary model. TreeTime uses the mugration model with and with-out sampling bias correction (shown in parentheses). SAASI uses the true parameter values that generated the tree. SAASI^*∗*^ uses estimated phylodynamic parameters but is provided with the true sampling rate. Trees were generated with speciation rates *λ*_1_ = 3 and *λ*_2_ = 1.5, extinction rates *µ*_1_ = 0.05 and *µ*_2_ = 0.1, transition rates *q*_12_ = *q*_21_ = 0.3, and sampling rates *ψ*_1_ = 0.25 and *ψ*_2_ = 1.0.

**Supplementary Fig. 9.**
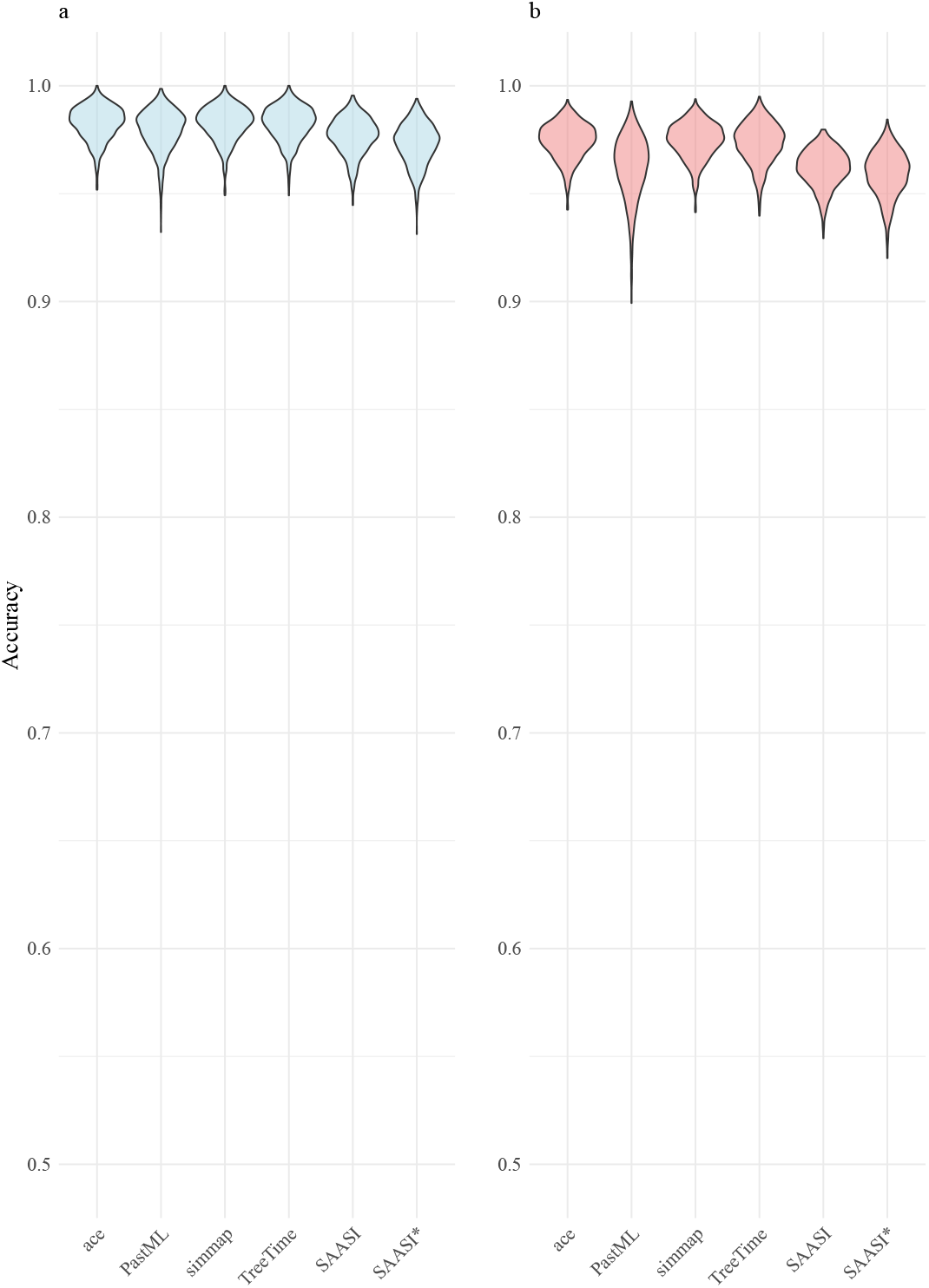
Impact of downsampling on ancestral state reconstruction accuracy under unbiased sampling (*ψ*_2_ = *ψ*_1_) and state dependent diversification across 1000 simulated trees. a: Consensus accuracy, defined as the fraction of correctly inferred ancestral node states. b: Probability accuracy, accounting for uncertainty in the node inference. Violin plots show the distribution of accuracy values across all simulations. ace and simmap use the equal-rates (ER) transition model. PastML employs the MPPA method with the F81 evolutionary model. TreeTime uses the mugration model with different sampling bias correction values (shown in parentheses). SAASI uses the true parameter values that generated the tree. SAASI^*∗*^ uses estimated speciation and extinction parameters but is provided with the true sampling rate. Trees were generated with speciation rates *λ*_1_ = 3 and *λ*_2_ = 1.5, extinction rates *µ*_1_ = 0.05 and *µ*_2_ = 0.1, transition rates *q*_12_ = *q*_21_ = 0.3, and sampling rates *ψ*_1_ = *ψ*_2_ = 0.5.

**Supplementary Table 1.** Summary statistics of accuracy comparisons under high sampling bias, with a mean tree size of 463 tips.

| Accuracy | Consensuses Accuracy |  | Probability Accuracy |  |
| --- | --- | --- | --- | --- |
|  | Mean | SD | Mean | SD |
| ace | 0.641 | 0.103 | 0.611 | 0.061 |
| PastML | 0.664 | 0.108 | 0.634 | 0.067 |
| simmap | 0.670 | 0.098 | 0.626 | 0.060 |
| TreeTime | 0.707 | 0.134 | 0.678 | 0.102 |
| TreeTime (correction) | 0.699 | 0.147 | 0.689 | 0.130 |
| SAASI | 0.934 | 0.021 | 0.912 | 0.019 |
| SAASI* | 0.920 | 0.023 | 0.818 | 0.037 |

**Supplementary Table 2.** Summary statistics of accuracy comparisons under moderate sampling bias, with a mean tree size of 2669 tips.

| Accuracy | Consensuses Accuracy |  | Probability Accuracy |  |
| --- | --- | --- | --- | --- |
|  | Mean | SD | Mean | SD |
| ace | 0.963 | 0.017 | 0.961 | 0.017 |
| PastML | 0.976 | 0.010 | 0.970 | 0.011 |
| simmap | 0.963 | 0.017 | 0.961 | 0.017 |
| TreeTime | 0.975 | 0.010 | 0.970 | 0.011 |
| TreeTime (correction) | 0.973 | 0.010 | 0.964 | 0.011 |
| SAASI | 0.976 | 0.007 | 0.964 | 0.008 |
| SAASI* | 0.975 | 0.008 | 0.968 | 0.010 |

**Supplementary Table 3.** Impact of downsampling on accuracy under high sampling bias: comparison between downsampled only, no downsampling, and downsampled with missing transitions datasets. The accuracies calculated in TreeTime, SAASI, and SAASI^*∗*^ only consider the tree under no downsampling.

| Accuracy | Subsampled tree only |  |  |  | With subsampled tree |  |  |  |
| --- | --- | --- | --- | --- | --- | --- | --- | --- |
|  | Mean Cons. Acc. | SD Cons. Acc. | Mean Prob. Acc. | SD Prob. Acc. | Mean Cons. Acc. | SD Cons. Acc. | Mean Prob. Acc. | SD Prob. Acc. |
| ace | 0.990 | 0.014 | 0.985 | 0.013 | 0.327 | 0.034 | 0.325 | 0.034 |
| PastML | 0.983 | 0.025 | 0.910 | 0.034 | 0.325 | 0.035 | 0.301 | 0.036 |
| simmap | 0.989 | 0.014 | 0.985 | 0.014 | 0.327 | 0.034 | 0.325 | 0.035 |
| TreeTime | 0.984 | 0.021 | 0.963 | 0.028 | 0.325 | 0.034 | 0.318 | 0.035 |
| TreeTime (correction) | 0.699 | 0.147 | 0.689 | 0.130 | - | - | - | - |
| SAASI | 0.934 | 0.021 | 0.912 | 0.019 | - | - | - | - |
| SAASI* | 0.920 | 0.023 | 0.818 | 0.037 | - | - | - | - |

**Supplementary Table 4.** Impact of downsampling on accuracy under moderate sampling bias: comparison between downsampled only, no downsampling, and downsampled with missing transitions datasets. The accuracies calculated in TreeTime, SAASI, and SAASI^*∗*^ only consider the tree under no downsampling.

| Accuracy | Subsampled tree only |  |  |  | With subsampled tree |  |  |  |
| --- | --- | --- | --- | --- | --- | --- | --- | --- |
|  | Mean Cons. Acc. | SD Cons. Acc. | Mean Prob. Acc. | SD Prob. Acc. | Mean Cons. Acc. | SD Cons. Acc. | Mean Prob. Acc. | SD Prob. Acc. |
| ace | 0.988 | 0.009 | 0.984 | 0.008 | 0.578 | 0.033 | 0.575 | 0.033 |
| PastML | 0.980 | 0.015 | 0.927 | 0.030 | 0.572 | 0.034 | 0.542 | 0.035 |
| simmap | 0.988 | 0.009 | 0.984 | 0.008 | 0.578 | 0.033 | 0.575 | 0.033 |
| TreeTime | 0.985 | 0.012 | 0.972 | 0.017 | 0.576 | 0.033 | 0.568 | 0.034 |
| TreeTime (correction) | 0.973 | 0.010 | 0.964 | 0.011 | - | - | - | - |
| SAASI | 0.976 | 0.007 | 0.964 | 0.008 | - | - | - | - |
| SAASI* | 0.975 | 0.008 | 0.968 | 0.010 | - | - | - | - |

**Supplementary Table 5.** Summary statistics of accuracy comparisons under uniform sampling, with a mean tree size of 1360.

| Accuracy | Consensuses Accuracy |  | Probability Accuracy |  |
| --- | --- | --- | --- | --- |
|  | Mean | SD | Mean | SD |
| ace | 0.983 | 0.008 | 0.974 | 0.008 |
| PastML | 0.979 | 0.010 | 0.962 | 0.014 |
| simmap | 0.983 | 0.008 | 0.975 | 0.001 |
| TreeTime | 0.982 | 0.008 | 0.973 | 0.010 |
| SAASI | 0.977 | 0.008 | 0.962 | 0.009 |
| SAASI* | 0.973 | 0.010 | 0.959 | 0.011 |

**Supplementary Fig. 10.**
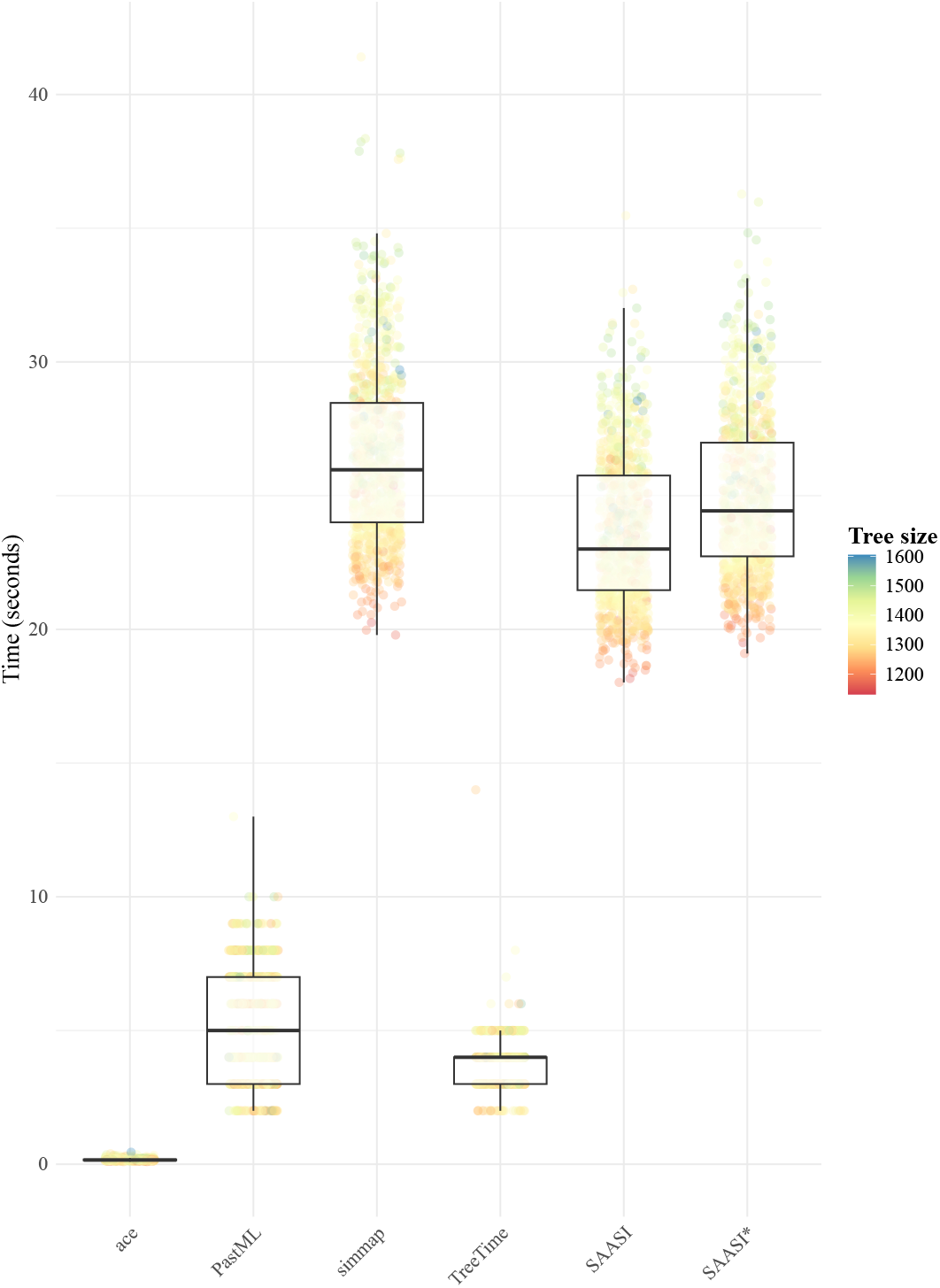
Computational time for ancestral state reconstruction methods. Running time is measured in seconds. Each point represents one simulation, with color indicating tree size (number of tips).

**Supplementary Fig. 11.**
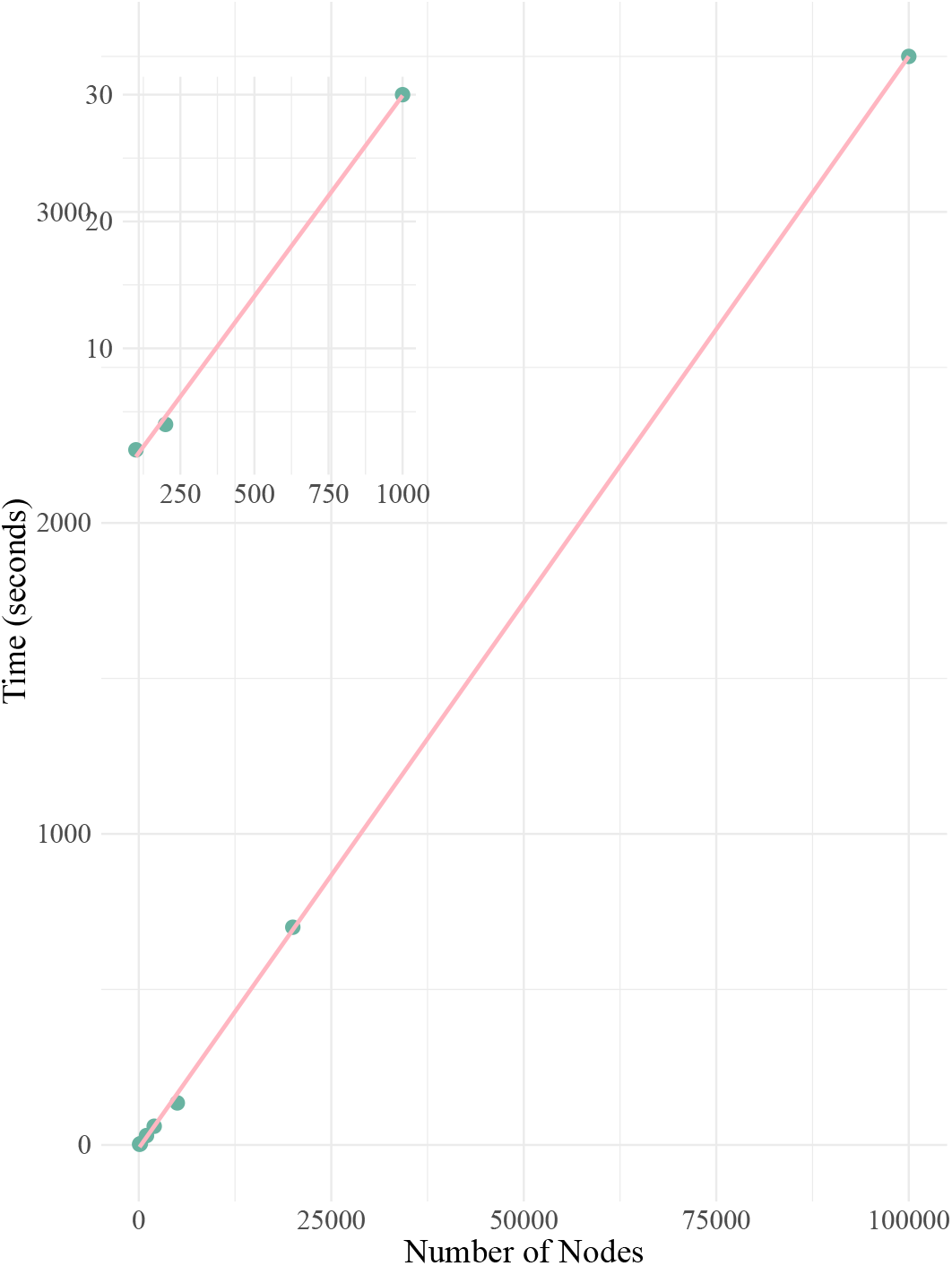
Runtime of SAASI across trees of different sizes. The x-axis represents the number of nodes, and the y-axis represents the times needed using SAASI. The red line shows the line of best fit. The small panel on the top left is a zoomed-in view of small tree sizes.

**Supplementary Fig. 12.**
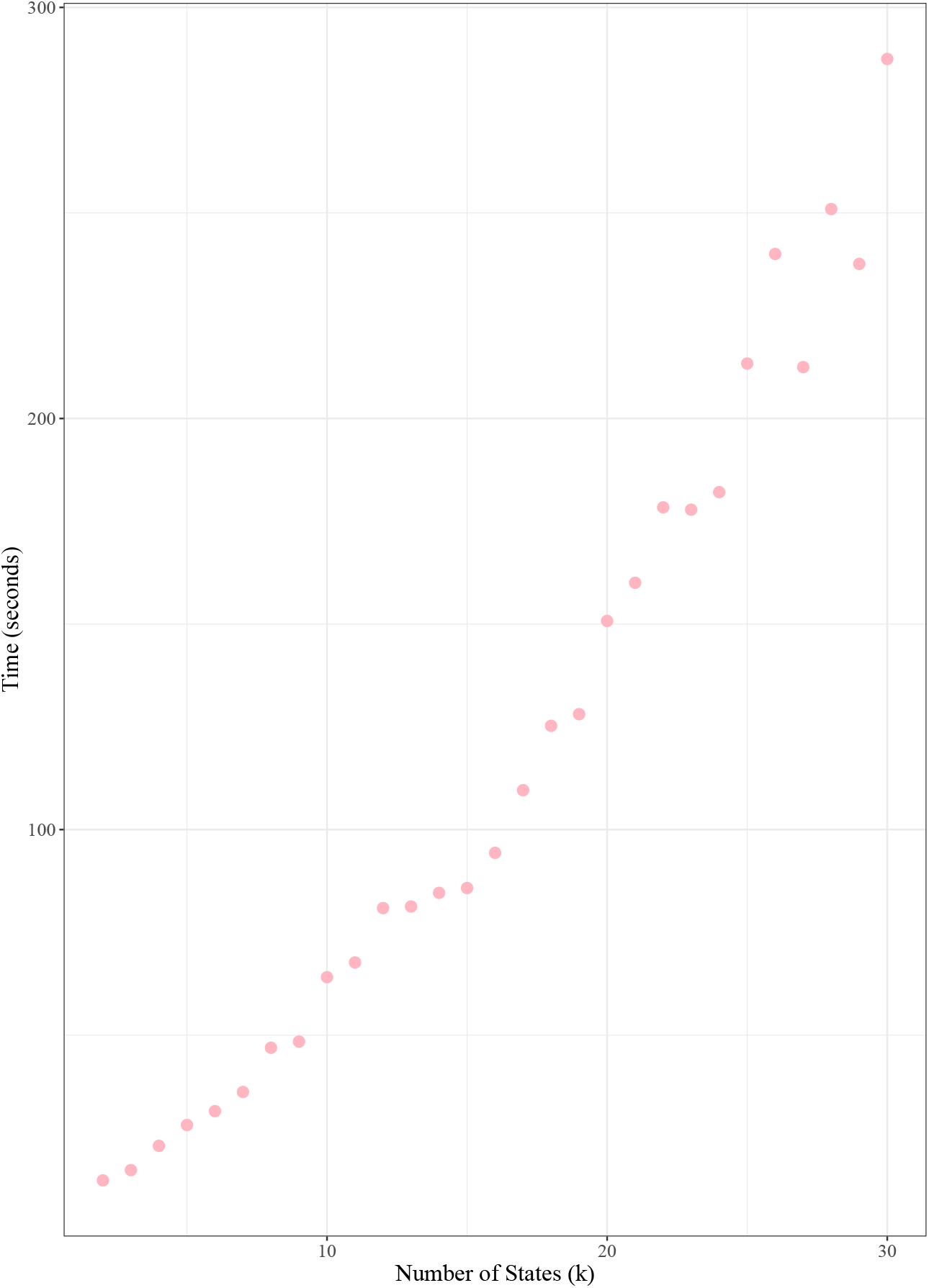
Runtime of SAASI across trees of different numbers of states. The x-axis represents the number of states, and the y-axis represents the times needed using SAASI.

